# Arrow heads at Obi-Rakhmat (Uzbekistan) 80 ka ago

**DOI:** 10.1101/2025.01.14.632810

**Authors:** Hugues Plisson, Alëna V. Kharevich, Vladimir M. Kharevich, Pavel V. Chistiakov, Lydia V. Zotkina, Malvina Baumann, Eric Pubert, Ksenya A. Kolobova, Andrei I. Krivoshapkin

**Affiliations:** PACEA UMR5199, Université de Bordeaux, France; Institut of Archaeology and Ethnography SB RAS Novosibirsk, Russian Federation; Traceolab, Université de Liège, Belgium; Zoostan, IRL 2033, CNRS & KazNU, Almatys, Kazakhstan; APSACA, National Center of Archeology, Tashkent, Uzbekistan

**Keywords:** Projectile head, bow and arrow, Central Asia, Uzbekistan, Middle Palaeolithic, Transition Middle to Upper Palaeolithic

## Abstract

When they are occasionally found in Middle Palaeolithic Neanderthal settlements, lithic weapons heads are large and do not differ in size, shaping or type from those used for activities other than hunting, such as plant gathering or butchery. The presence in a same assemblage of various types of armatures, some of which are microlithic and designed for this purpose, has only been documented in Modern Humans sites.

Recent studies show that light projectile points, which were to become the structuring element of Upper Palaeolithic lithic industries, were already present in its formative stages. However, they remain marginal in debates regarding the Middle to Upper Paleolithic transition.

We present here the first results of a search for weapon heads in the oldest layers of the Obi-Rakhmat rock shelter, at around 80 ka. This site, located in the western foothills of the Tian Shan Mountains, northeastern Uzbekistan, has yielded a blade-based lithic industry which is forming part of the continuity of the Levantine Early Middle Paleolithic but with several innovative traits. The assemblage is based on a systematic blade production (regular thick narrow blades from unipolar and bipolar sub-prismatic and narrow-faced cores, thin and wide blades from flat-faced Levallois-like cores) along with shorter pieces from convergent or centripetal Levallois cores, and bladelets from burin-cores and other small cores.

Three types of projectile armature are identified from their impact marks: retouched points, bladelets and raw micropoints which were so far unseen due to their fragmentary state. The micropoints are too narrow for having been fitted to anything other than arrow shafts. They are similar to the armatures described in a pioneer settlement by Sapiens in the Rhône Valley, France, 25,000 years later.

## Introduction

### Archeological perspective

Instrumented hunting is a distinctive trait of the Homo genus. Due to the impact of meat consumption on the hominisation process, both cognitively and behaviourally [1], the search for archaeological evidence of the weapons used in the distant past is of prime importance with a particular attention paid to the oldest occurrences.

We present here the first results of a search for weapon points in the oldest levels of the Obi-Rakhmat rock shelter in Uzbekistan at around 80 ka. The lithic industry of this settlement is forming part of the continuity of the Levantine Early Middle Paleolithic but with several innovative traits [2].

Increasing studies show that middle or small sized lithic points which are part of the typological characterisation of the Initial or Early Upper Palaeolithic assemblages were projectile heads [3–6], probably mechanically delivered [7–9]. They mark a technical break with the Middle Palaeolithic; from then on, projectile armatures will become the central structuring element of lithic industries (Bordes and Teyssandier, 2011). In spite of this, they remain marginal in the debates regarding the Middle-to Upper Palaeolithic *transition*. Already suspected of being present in Obi-Rakhmat sequence despite its age [10], light projectile points deserve attention.

### Weapons in focus

Various criteria have been used to recognise prehistoric hunting weapons. The first is analogy with the shapes of comparable objects known from ethnographic records or from modern sporting or play practices. This is the case with javelins and throwing sticks from the ancient Palaeolithic [11]. In a more sophisticated approach, these are various acuteness indices calculated for lithic points on ethnographic and experimental bases, such as TCSA and TCSP [12–17]. But these are only theoretical potentialities [18,19]. As F. Bordes wrote: “*It could just as easily be argued that the sockets of bronze spearheads were used to cut rounds out of pie dough*” [« On pourrait tout aussi bien soutenir que les douilles des pointes de lance en bronze servaient à découper des ronds dans la pâte à tarte » 20]. What’s more, these indices don’t distinguish between simply tapered heads and those with cutting capacity, which makes a significant difference in real-life hunting, and they don’t take into account the incidence on the penetration of the hafting device which depends on the morphology of the basal part. A more reliable basis is provided by functional clues. The most obvious of these is when the tip of a perforating projectile is found stuck in a bone, but such find is infrequent [21–29]. It is much more common to find impact damage on the lithic or bone points. However, these are variable, depending on the projectile design, the ballistic parameters and the impacted target. In fact, unlike the tools used for cutting, scraping, drilling, etc., where the cause-effect relationship between wear and use is direct and invariant, since the same causes produce always the same effects, the identification of a projectile point or insert is an extrapolation based on the direction of the violent stress that broke the artefact. Causes other than being at the tip of a spear or arrow can lead to axial compressive stresses: knapping accident [30–32] (S1 - 2 Fig), certain types of shaping [33], hard butchering, use as a chisel, accidental dropping, etc. In some cases, the distinction is easy to make at the scale of the artefact itself, in others it is less so if a range of criteria is not taken into account. A good example is provided by obsidian points from the Ethiopian rift dated to 279 ka years ago and interpreted as javelin tips on the basis of their shape, apical removals and velocity-dependent microfracture features [34]. This last innovative criterion is physically relevant, but may be not the presumed cause of the energy involved. A larger study on similar assemblages, based on a technological analysis, suggests that the recurrent apical removals on such pointed artifacts result from a rejuvenation by the lateral tranchet blow technique [35]. More commonly confusing are the minor damages that often occur at the tip of the pointed tools, their most exposed and fragile part. The negative of a tiny burin spall can turn a Levallois triangular flake into a projectile head [36, fig. 5].

There are two levels of analysis in the identification of a projectile armature. The first focuses on the impact traces, according to the following criteria:

- the morphology of the damage, which reflects the orientation of the stress [37].

- the extent of the damage, which depends on the energy dissipated on impact (resulting from the momentum of the projectile [« Momentum determines THE AMOUNT OF FORCE which an arrow has available to it for penetration » [38]], the hardness of the impacted material, the impact angle, and the toughness of the attachment of the point to the shaft), according to the material, shape and dimensions of the point or insert.

- the variability of the damage (type, dimension, combination) within the set of armatures, which reflects the variability of the shots.

- the location of the damage, depending on the position on the shaft and attachment.

The second level of analysis relates to interdependent functional parameters concerning the manufacture of the weapon, its ballistic properties and its lethal potential. What these parameters have in common is that they result in a morphological and dimensional standardization.

On the one hand, for thrusted spears the constraints are at the level of the junction between the armature and the shaft and its expected robustness [39]; on the other, for the projectiles, they are at the level of the ballistic precision and the energy involved in the shot [40].

In the high-risk, high-return activity that hunting typically is, any reduction in uncertainty is essential. On the technical side, this means a design that preserves the effectiveness of thrusting weapons or that ensures reproducibility of effective shots, which are both linked to strict compliance with physical principles that leave little room for improvisation [40,41]. In this complex system, the nature of the game, its ethology, reactivity, aggressiveness and resilience are also part of the parameters.

To put it simply, the design of a point will be very different depending on whether you’re hunting rabbits with a bow or aurochs with a spear.

### Obi-Rakhmat

The Obi-Rakhmat rock-shelter is located in the Paltau valley, at the south-western end of the Talassky Alatau range of the Tien Shan mountains, in northeastern Uzbekistan, 100 km of Tashkent. (N41°34’08.8” and E70°08’00.3”) (Fig 1). Carved into the Paleozoic limestone at an altitude of 1,250 m, it takes the form of a niche measuring 20 m in width and 9 m in length, with a southern orientation. Since the 1960s, several excavation campaigns have been conducted, initially by the Institute of History and Archaeology of the Uzbek Academy of Sciences [42], and later in collaboration with the Institute of Archaeology and Ethnography, Siberian Branch of the Russian Academy of Sciences [43,44].

**Fig 1.**
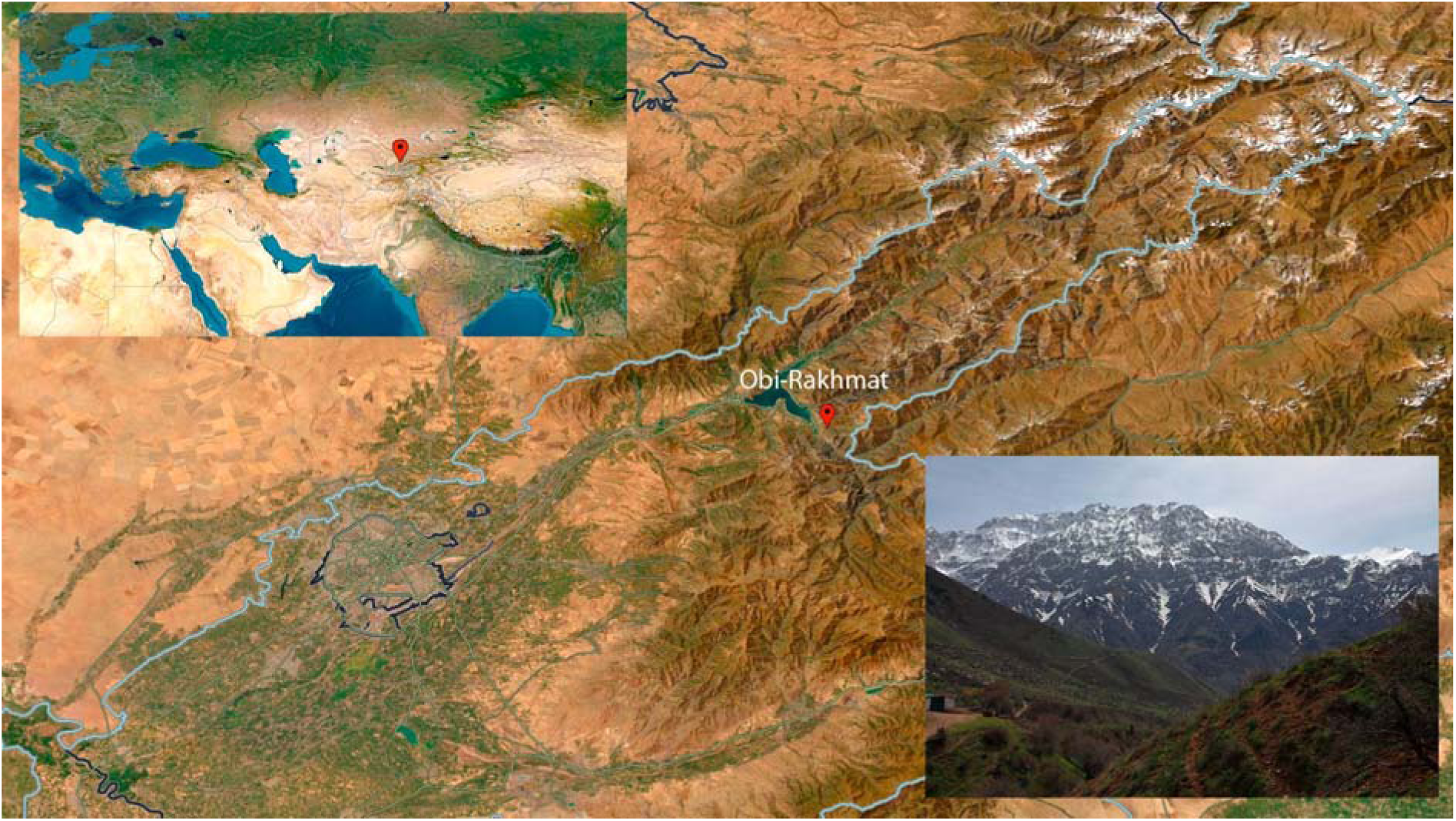
Location of Obi-Rakhmat - N41°34’08.8” and E70°08’00.3”, 1,250m asl - (maps courtesy of NASA) and view from the site.

The exposed stratigraphy (Fig 2) consists of 21 sedimentary levels spanning a depth of 10m [45], all of which contain archaeological material. The lithic industry, made from local silicified limestone, is homogeneous and characterized by the production of large blades from unipolar or bipolar and narrow-faced cores and bladelets from core-burins and bladelet cores of various morphology. These reduction strategies coexist alongside Levallois concept which is manifested by the presence of convergent or centripetal Levallois cores and flat-faced cores (Levallois-like). The typological tools include blades—often pointed and/or retouched— splintered pieces, burins, end- and side scrapers, denticulated, borers and retouched flakes. Notably, Levallois (mostly elongated) and Mousterian points are also present, the morphology of which is various and corresponds to the typology of retouched points in the Levantine Early Middle Palaeolithic [46,47]. This composition has led to comparisons between the Obi-Rakhmat lithic assemblage and the Late Middle Palaeolithic and Early Upper Palaeolithic blade industries [44,48] from Southwest Asia (Levant and neighboring territories; Bar-Yosef and Kuhn, 1999; Boaretto et al., 2021; Meignen, 2007, 2000) and the Siberian Altai [53–56]. Attempts to date the site have yielded heterogeneous results, as is often the case when applying different methods (^14^C AMS, U-series, ESR). However, they fix a chronological range from 90 ka for the deepest strata to 40 ka for the uppermost levels [57–59].

**Fig 2.**
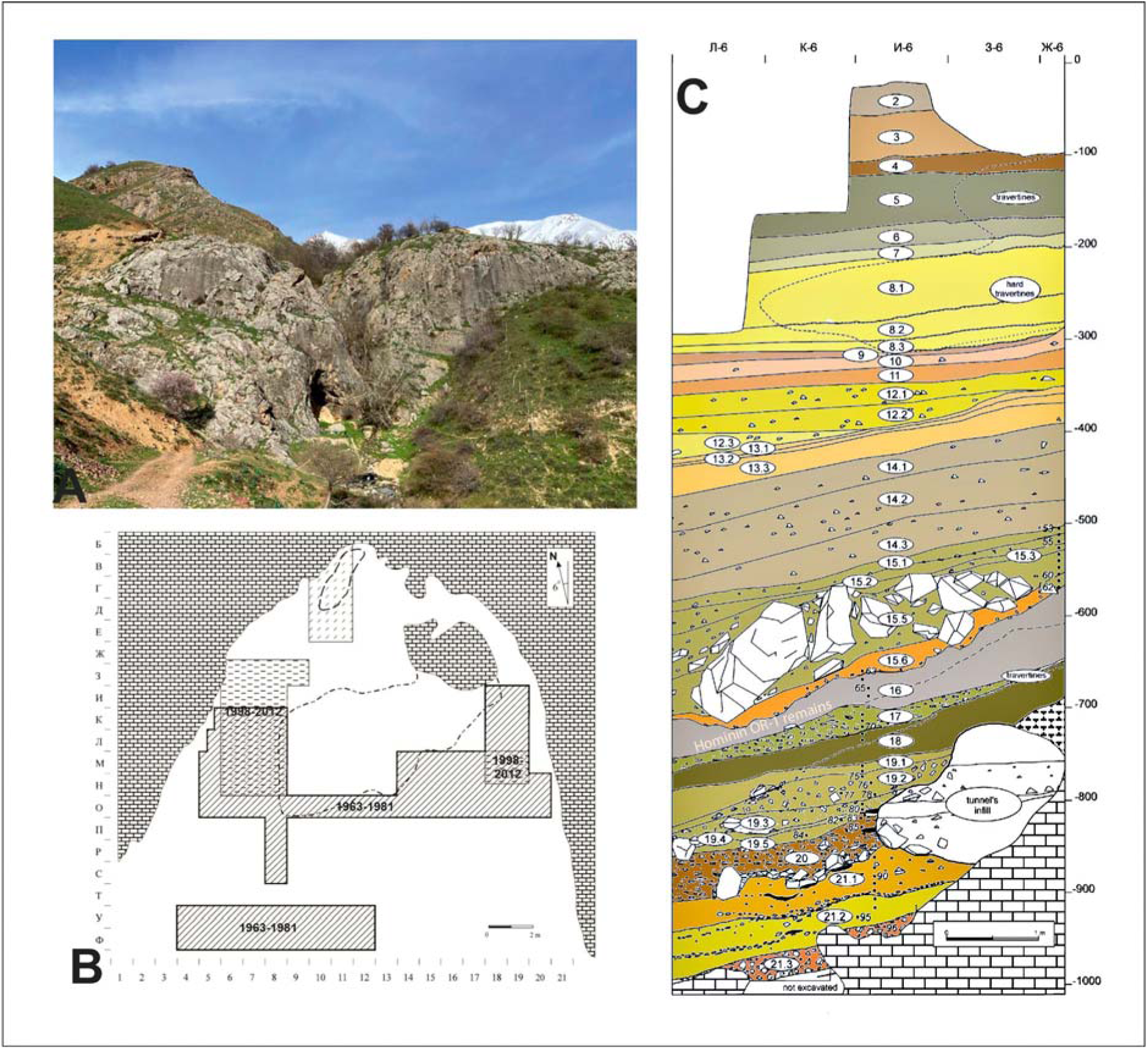
Stratigraphy of Obi-Rakhmat rock shelter and map of the excavation.

The faunal spectrum at Obi-Rakhmat is limited and shows little variation throughout the stratigraphy (Fig. 2, B). It is dominated by goat (*Capra sibirica*) and red deer (*Cervus elephus*), with smaller contributions from wild boar (*Sus scofra*), roe deer (*Capreolus capreolus*), golden jackal (*Canis aureus*), fox (*Vulpes vulpes*), marmot (*Marmota sp.*) and hare (*Lepus sp*.). This assemblage reflects a combination of steppe and forest environments [60,61], consistent with palynological data [62]. Remains of large carnivores, including cave lions, hyenas and bears, are rare. In 2003, human remains were discovered at the site: 6 left maxillary teeth and 121 skull fragments from a single juvenile individual (9-12 years old). While the dental morphology aligns more closely with Neanderthal populations, the mosaic of cranial morphological features prevents a definitive attribution, leaving open a classification as an archaic *Homo sapiens* [48,63–65]. Finally, elements of bone industry have been identified among the faunal remains [66,67].

## Material and methods

The analysed sample comes from the collection of the 2001 excavation campaign led by A. Krivoshapkin which is currently under study. It includes typological pieces and a search for triangular shapes among the bags of lithic debris from layers 20 to 21 stored at the National Center of Archeology in Tashkent, Uzbekistan. The first sorting was done with the naked eye, then the selected pieces were examined with a stereoscopic microscope (Wild M1B / x7, x14 magnification) before those with the least eroded surfaces were analysed with a reflective optical microscope (Olympus BHM, bright field with DIC / x50, x100, x200, x500 magnification) in search for microscopic linear impact traces - MLIT [68].

Photomacrographs of the impact traces were captured in raw format using a Canon EOS 60D camera equipped with a Canon EF-S 60 mm f/2,8 Macro USM lens and photomicrographs using a Nikon D750 on the phototube of the microscope. Multi-focus shots were processed with Helicon Software.

The artefacts were scanned using a Solutionix D500 3D scanner. The 3D PDFs of the supplementary data were produced in Acrobat Pro 9 from the STL files converted in MeshLab 2021.10.

The macroscopic diagnostic features were the morphology of the fractures and of the lateral damage as described in 40 years of publications [3,4,37,eg. 69–73]. Our personal experience is based on a corpus of over 500 flint points and barbs of various shapes used as arrow, dart and spear heads (Fig 3) on medium and big sized mammals [74–80].

**Fig 3.**
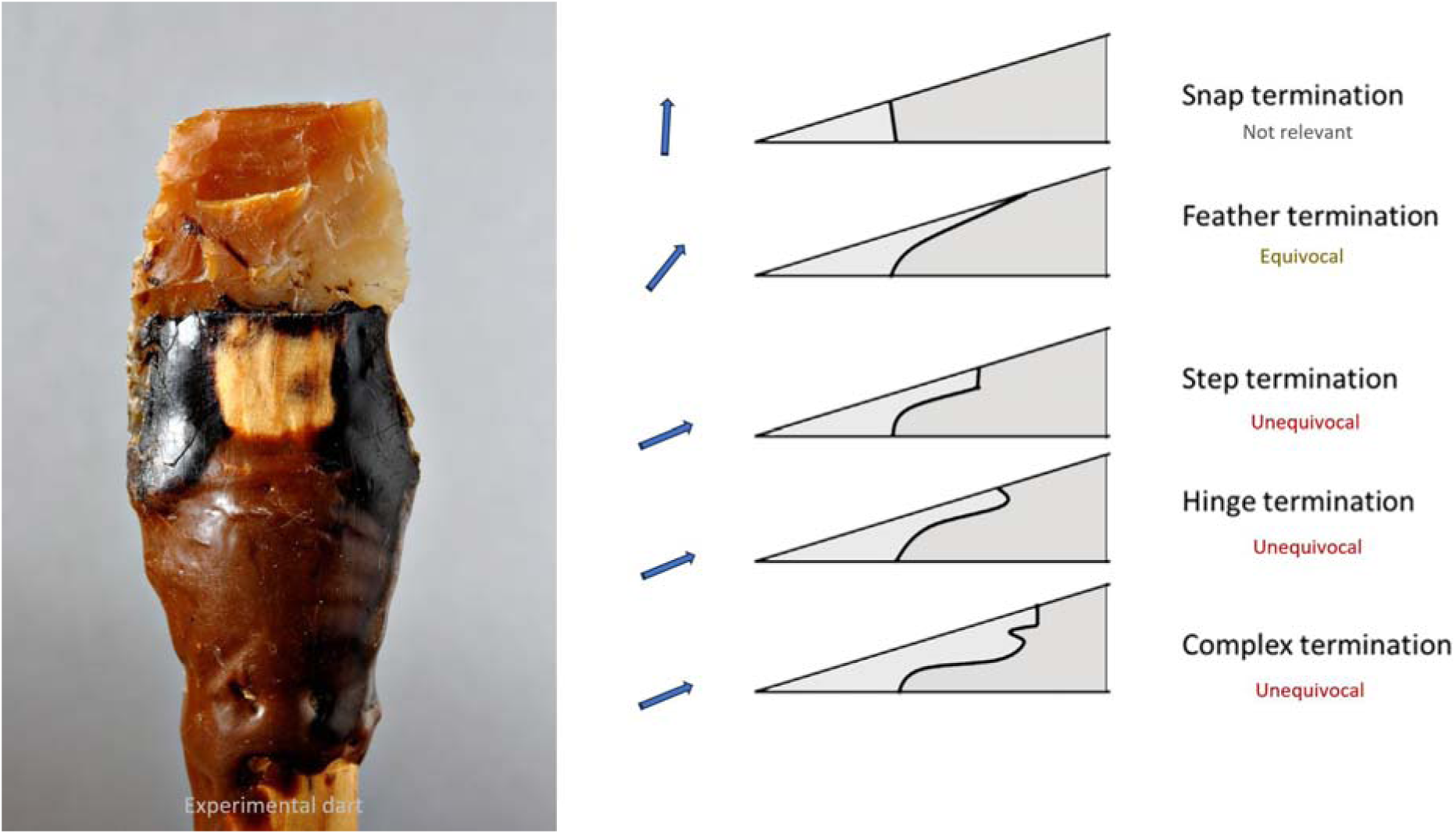
Terminations of bending fractures. Left: experimental Solutrean leaf point used as dart head, broken at impact, with long step-terminating bending fracture (TFPS Collective Research program). Right: profile of the different types of termination according to the orientation of the stress [37,81]

Additional preliminary experiments (S21 Fig) were undertaken with the local silicified limestone to test the debitage scheme assumed on the basis of the shape of the micropoints and potential cores seen in the lithic assemblage. This step enabled us to observe the knapping accidents that could mimic impact damage. Then 12 of the micropoints thus produced were fixed to 8mm-diameter commercial wooden arrows shafts using a mixture 80/20% of bitumen (natural outcrops exist in the region) and pulverized charcoal, without binding (this would be a technical absurdity on continuous cutting edges) for being shot with a modern laminated bow (36 lbs) on a complete uneviscerated carcass of a small ungulate hung in anatomical position in order to start documenting impact breakage and microscopic linear impact traces for the shape and raw material under study.

In order to reconstruct the technologies of point and micro-point production, an attributive analysis of cores and truncated-faceted pieces, as well as points and micro-points with use wear traces was carried out. The reconstruction of reduction sequences of the cores, the micro-cores and the truncated-faceted pieces was achieved through scar-pattern analysis. Comparative studies were carried out using a statistical approach, including Mann–Whitney pairwise test.

## Results

### Traceological inventory

On the basis of the macroscopic criteria usually used in publications and of our own experimental corpus, we have selected 20 pieces which can be regarded as projectile armatures from the inventory in progress of the lithic industry from the deepest levels (20-21) of Obi-Rakhmat. Most of the artefacts, due to the raw material and surface alteration are unsuitable for microscopic analysis, however MLIT have been found on two micropoints. Consequently, the determinations are based on the morphology of the fractures, which depends on the proportion of compressive and bending stress that caused the material to break by buckling or percussion (Fig 3) [37]. To enable each specialist to make their own judgement, the 3D model of each piece is provided in supplementary information.

The sample is still small, but it comes from an excavated area of 20 square meters and already shows a diversity of distinctive traces and morphological recurrences.

The armatures are of 3 types: more or less massive points (around 30 grams for nearly complete specimens), micropoints (between 1 and 2 grams) and bladelets (Fig 4).

**Fig 4.**
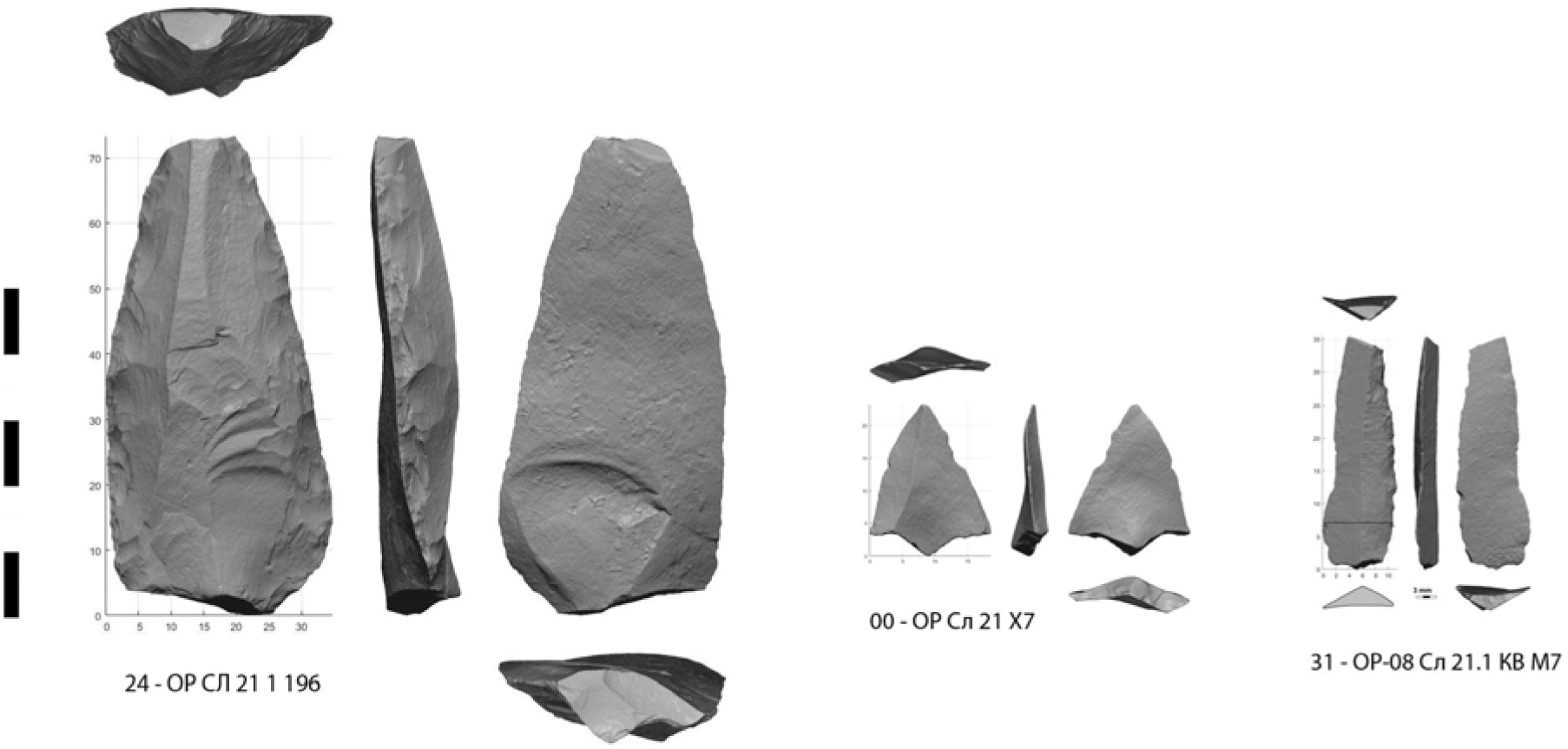
Three types of lithic weapon armature identified in layers 20 and 21 of Obi-Rakhmat: Medium sized retouched point, micropoint and bladelet.

The massive points (Fig 5) are illustrated by 2 almost complete retouched points (Fig 5: 28, S1 Fig, S1 File, S2 File, S2 File) (38 mm and 41 mm wide) and an apical half (Fig 5: 22, S3 Fig, S3 File), all 3 crushed at the tip by an axial impact. An apical fragment of retouched point from layer 19 is similar, having been badly chipped by a tangential impact to its right edge along the same axis (Fig 5: 104, S4 Fig, S4 File). The proximal thinning on both sides of the entire samples suggests a hafting layout. A thicker but more elongated point (Fig 5: 24, S5 Fig, S5 File) (30 mm wide) has an apical fracture with a wavy feather termination on the lower side from the same kind of compressive stress.

**Fig 5.**
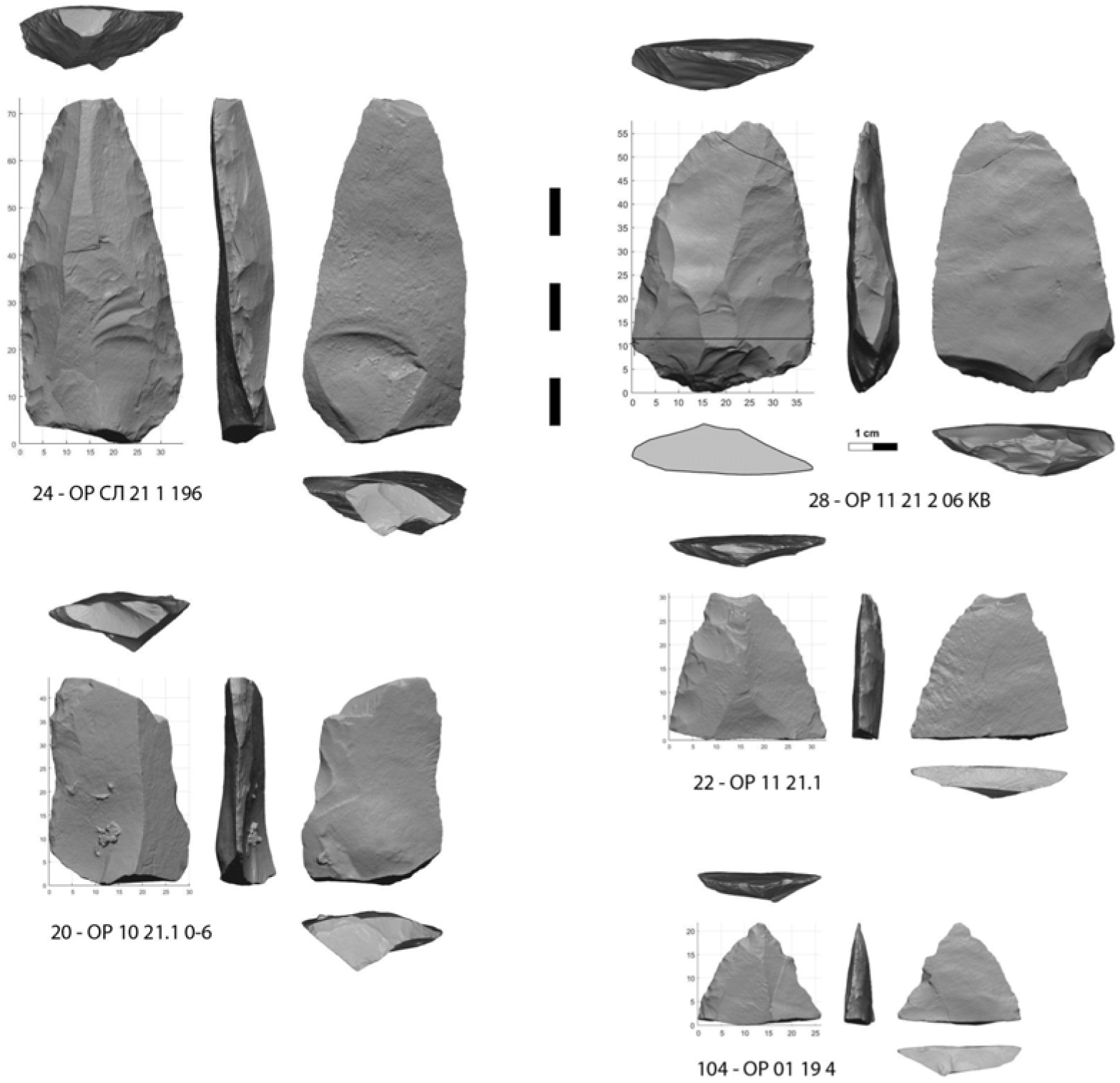
Medium sized retouched points crushed or broken at impact.

One large proximal fragment (Fig 5: 20, S6 Fig, S6 File) which could be from the same type of point than the previous one, also have a fracture attributable to its use as the tip of a spear, but lateral, which can only occur under the leverage of a long shaft.

Two apical fragments from more slender retouched points complete the set. One has a bending fracture with an atypical feather termination but here slightly twisted (S7 Fig, S7 File), while the other has a long spin-off fracture (S8 Fig, S8 File).

The second category is made up of 9 unretouched micropoints and 1 retouched micropoint (Fig 6, S9-16 Fig, S9-16 File), produced in a variety of ways, from simple triangular flakes to typical Levallois point. Two samples have both fracture from longitudinal stress and MLIT (Fig 7-8). The average width is 18.2 mm (minimum 15,3 – maximum 23,7 mm). The average weight of broken specimens is 1.4 grams (minimum 0.7 g - maximum 2.5 g). An unbroken Levallois micropoint (Fig 6: 00, S17 Fig, S17 File) measures 21.8 mm long x 17.7 mm wide, weighing 1.1 grams ; its edge damage is atypical (trampling or crushing). There is no basal shaping despite prominent bulbs. The location on the lower side of the fracture hinge for 6 specimens, half of which have elongated endings, whereas a flat surface is not conducive to their extension, suggests a mounting on the shafts that does not compensate for the prominence of the bulb, i.e. a mounting that is not perfectly in line with the axis (what is not optimal for the wake drag coefficient). The shafts were probably made from wood only, since no potential intermediate nor apical bone elements were found.

**Fig 6.**
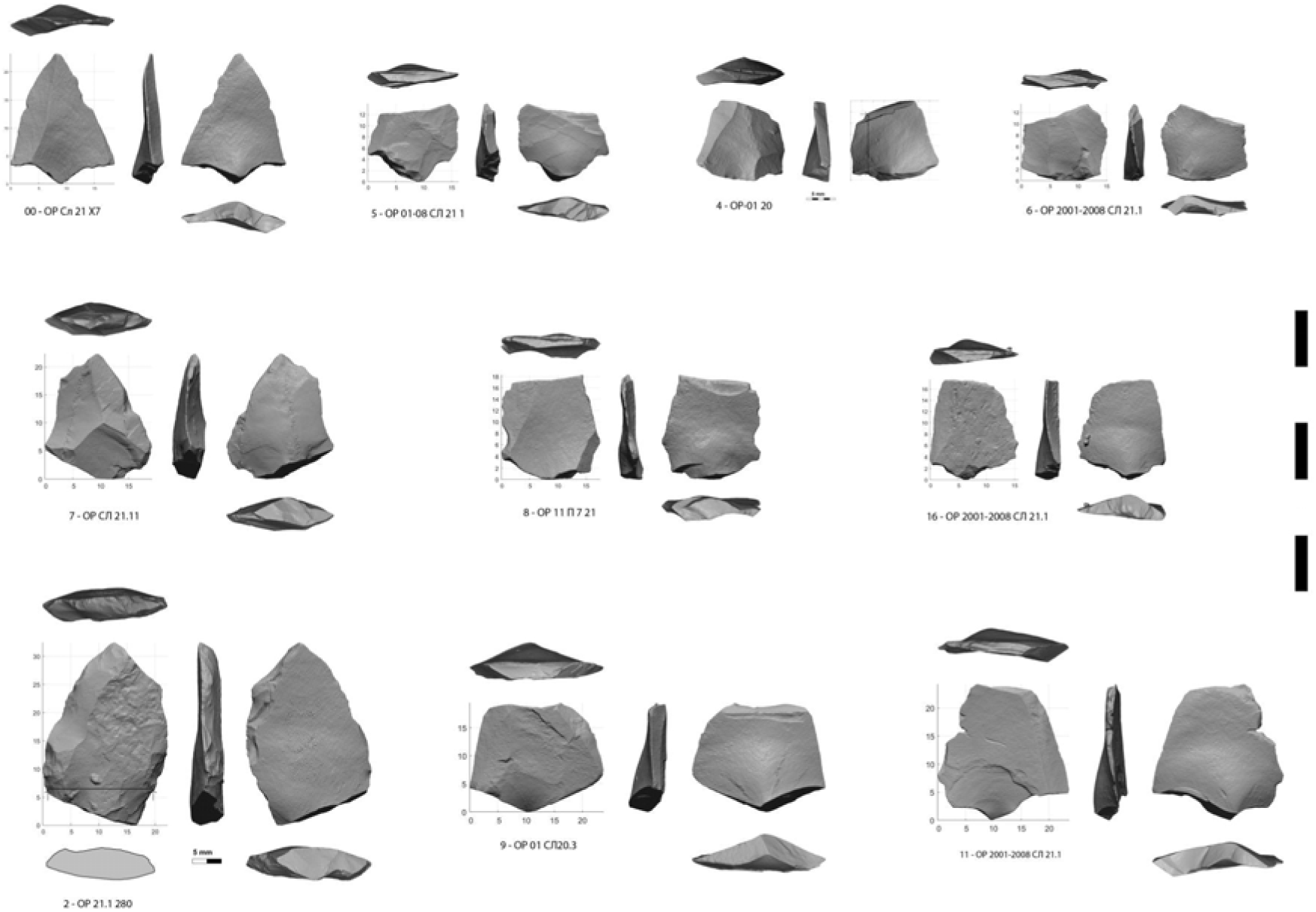
Micropoints broken or crushed at impact.

**Fig 7.**
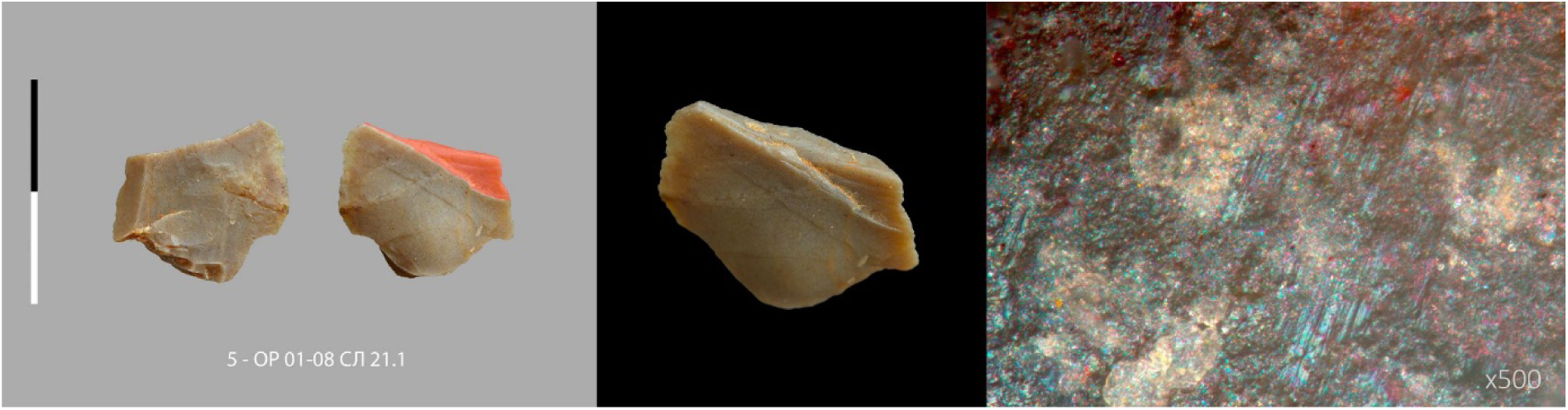
Micropoint n° 7 (5 - ОР 01-08 СЛ 21.1) with macroscopic and microscopic impact traces (MLIT). Centimetric scale.

**Fig 8.**
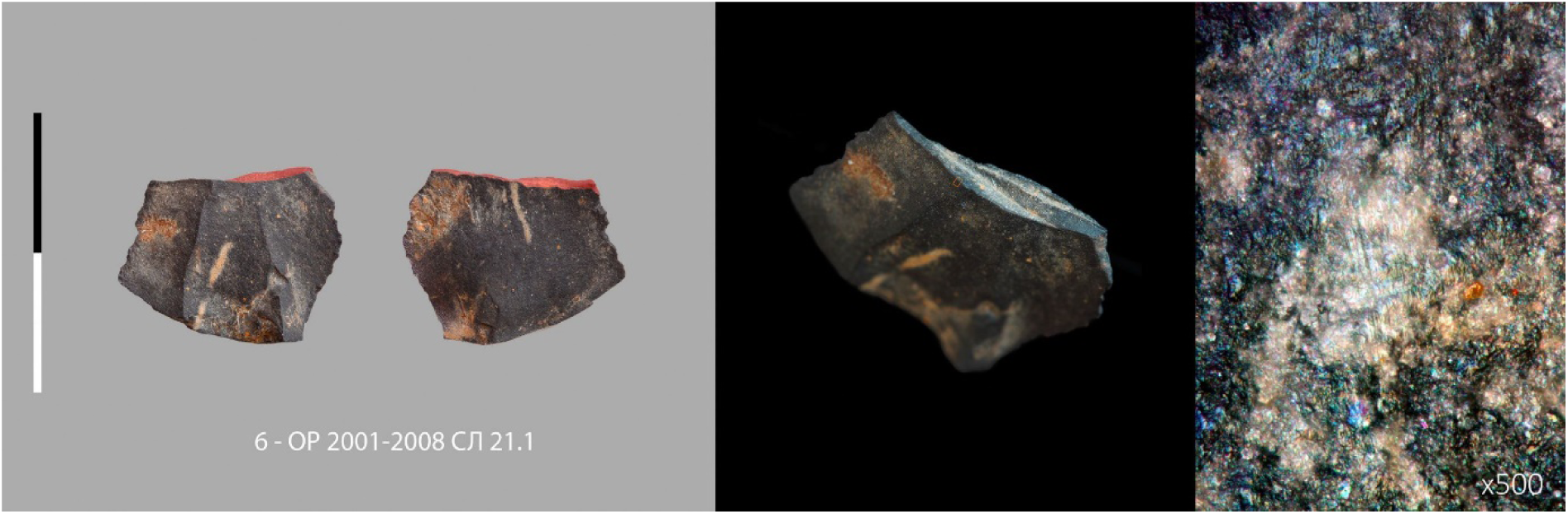
Micropoint n° 8 (6 - ОР 2001-2008 СЛ 21.1) with macroscopic and microscopic impact traces (MLIT). Centimetric scale.

The 3rd category (Fig 9) is the least represented here, as it was not included in our sorting criteria, which focused on axial armatures. Four incidental findings come from this sorting. They are raw bladelets. It is difficult to distinguish between those accidentally broken during knapping [30,32] and those broken by an axial use. Two were selected on the basis of the following criteria: one (Fig 9: 19, S18 Fig, S18 File), whose breakage by bending is equivocal, has a very discreet retouch along one edge produced by pressure, the second (Fig 9: 15, S19 Fig, S19 File), burnt, has been crushed by a tangential contact with a hard material, which is typical of projectile lateral inserts. A more discrete crushing is also observed on a genuine backed bladelet (Fig 9: 31, S20 Fig, S20 File).

**Fig 9.**
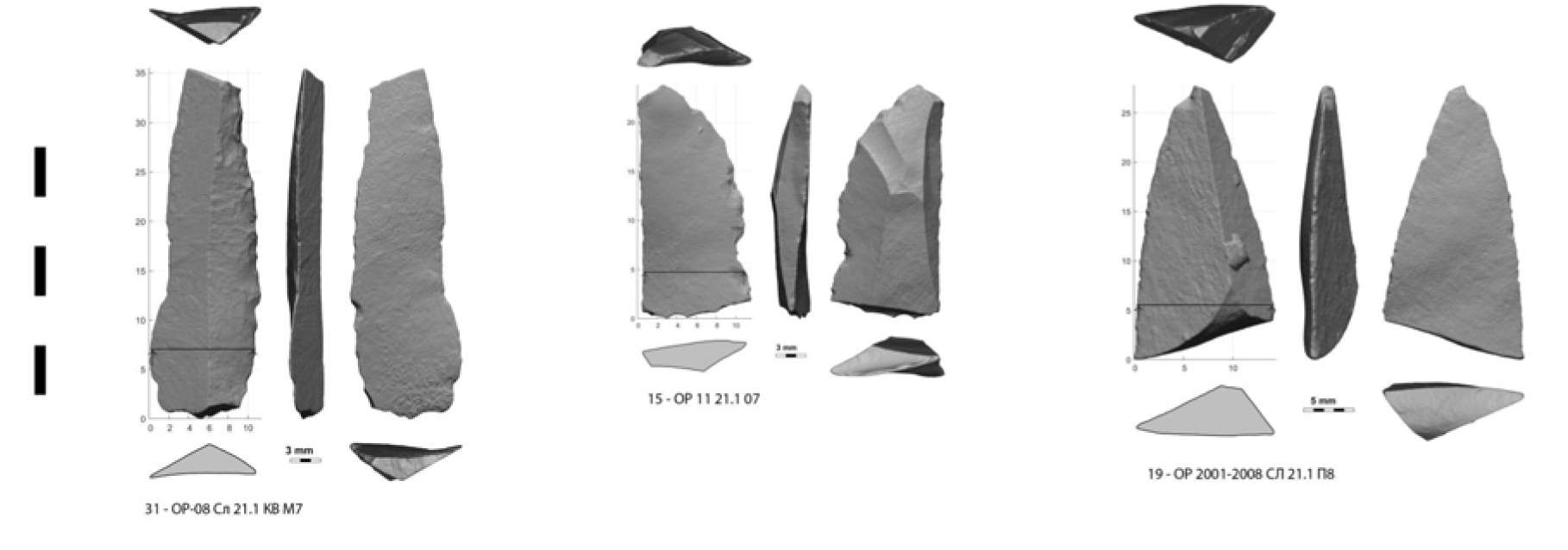
Bladelets crushed of broken at impact.

The set is very small, but sufficient to indicate the use of bladelets in the design of hunting weapons at Obi-Rakhmat, 80 ka ago. We could imagine that they complemented the micropoints, to extend the length of cutting edge, being thus part of a more robust projectile head compatible with dart foreshafts, but nothing in the design of the micropoints would ensure a continuity to prevent the bladelets from being pulled out at the penetration. More recent archaeological examples show the extreme attention paid to the continuity of lithic inserts [82]. A larger sample and therefore specific research would be needed to understand the function(s) of the bladelets, which are a remarkable element of the lithic production at Obi-Rakhmat.

### Technological characterization of points production

#### Point production

The Obi-Rakhmat assemblage contains a significant number of pointed blades and points (Fig 10). These were obtained both as part of the dedicated Levallois reduction sequences and as part of the blade core reduction (Fig 10).

**Fig 10.**
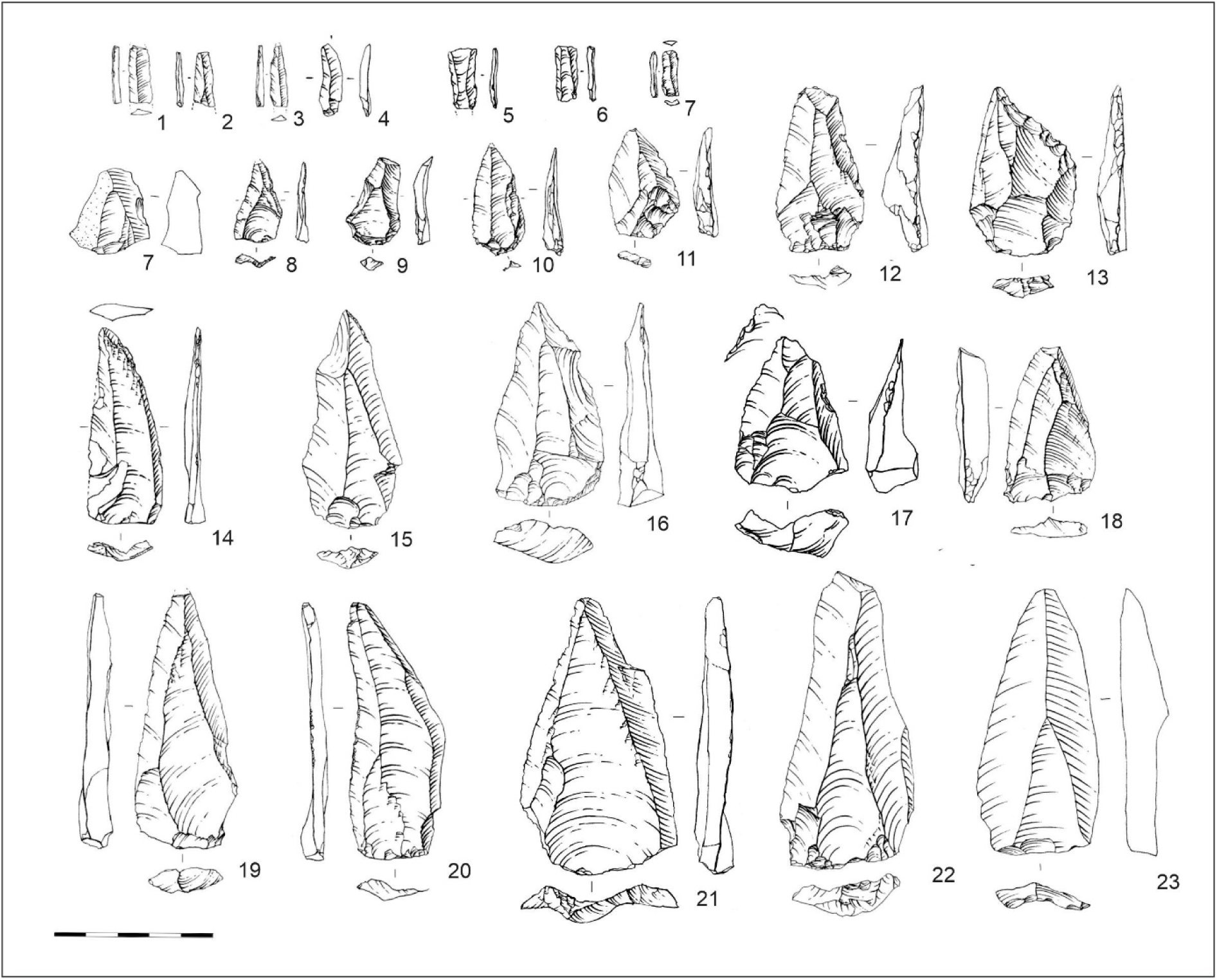
Convergent spalls from layers 20-21. 1-6 Bladelets; 7-10, 17-18, 21-23 Levallois points; 11-13, 16 Axial points; 14-15, 19-20 Elongated points.

In the assemblage of layers 20-21, the proportion of Levallois point cores (Fig 11 A) ranges from 5 to 30% (S1 table). According to scar-pattern analysis, pointed negatives are often recorded on blade cores as well. The collection includes flat-faced unidirectional (Fig 11, 2) and bidirectional (Fig 11, 3) cores, as well as a considerable proportion of narrow-faced cores (Fig 11, 4-5), which frequently exhibit pointed negatives. Another technological method (Fig 11 D), which is extensively represented in the collection, is associated with subprismatic unidirectional (Fig 11, 6) and bidirectional (Fig 11, 7) cores. Among the subprismatic cores, there are asymmetrical (semi-rotated) cores (Fig 11, 8) in which both large and narrow core faces are used together.

**Fig 11.**
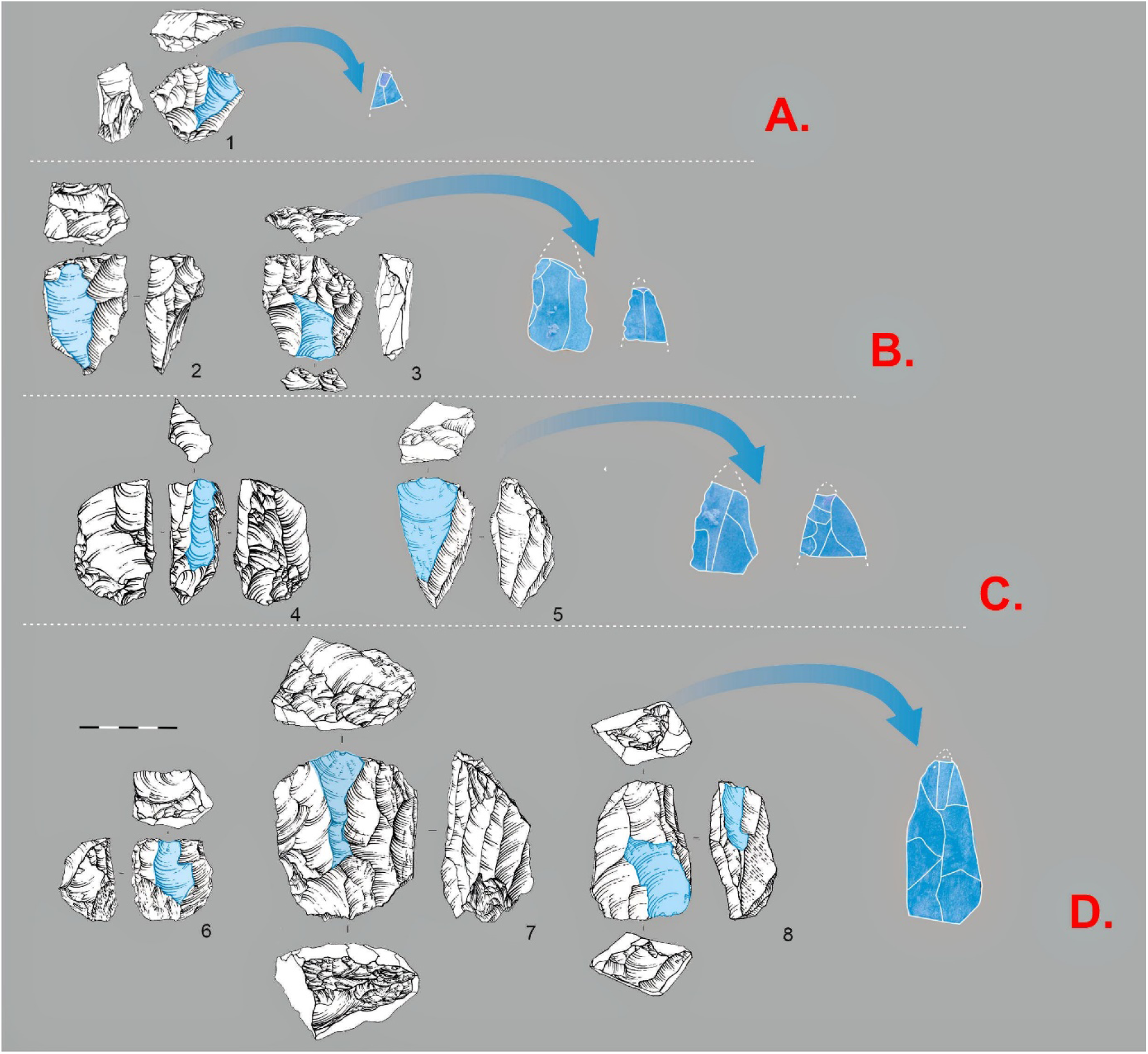
Technological chains of point and elongated blade production. A. 1 Levallois point core. B Flat faced cores: 2 Flat faced, unidirectional; 3 Flat faced, bidirectional. C. Narrow-faced cores: 4-5 Narrow-faced unidirectional. D Sub-prismatic cores: 6 Sub-prismatic unidirectional; 7 Sub-prismatic bidirectional; 8 Sub-prismatic asymmetrical (*Semi-tourné*).

#### Micropoint and bladelet production

On initial observation, it is challenging to identify a distinct chain within the Obi-Rakhmat assemblage that is specifically designed to produce micro-points. At the same time, a developed bladelet production is widely represented in the lithic industry (S1 table). The assemblage of lower layers is comprised of 20-30% bladelets (S1 table), with the proportion of bladelet cores reaching 50% (S1 table). A scar-pattern analysis of bladelet cores allowed us to record a variety pointed negatives on them (Fig 12). The bladelet cores used in the Obi-Rakhmat industry exhibit a high degree of diversity, with the presence of carenated (Fig 12, E), narrow-faced (Fig 12, A), burin-cores (Fig 12, B), flat-faced (Fig 12, D), and sub-prismatic (Fig 12, C) cores. The utilization of all these technological chains yielded micro points, as well as pointed bladelets that exhibit a morphological resemblance to the archaeological specimens that were selected as projectile armatures.

**Fig 12.**
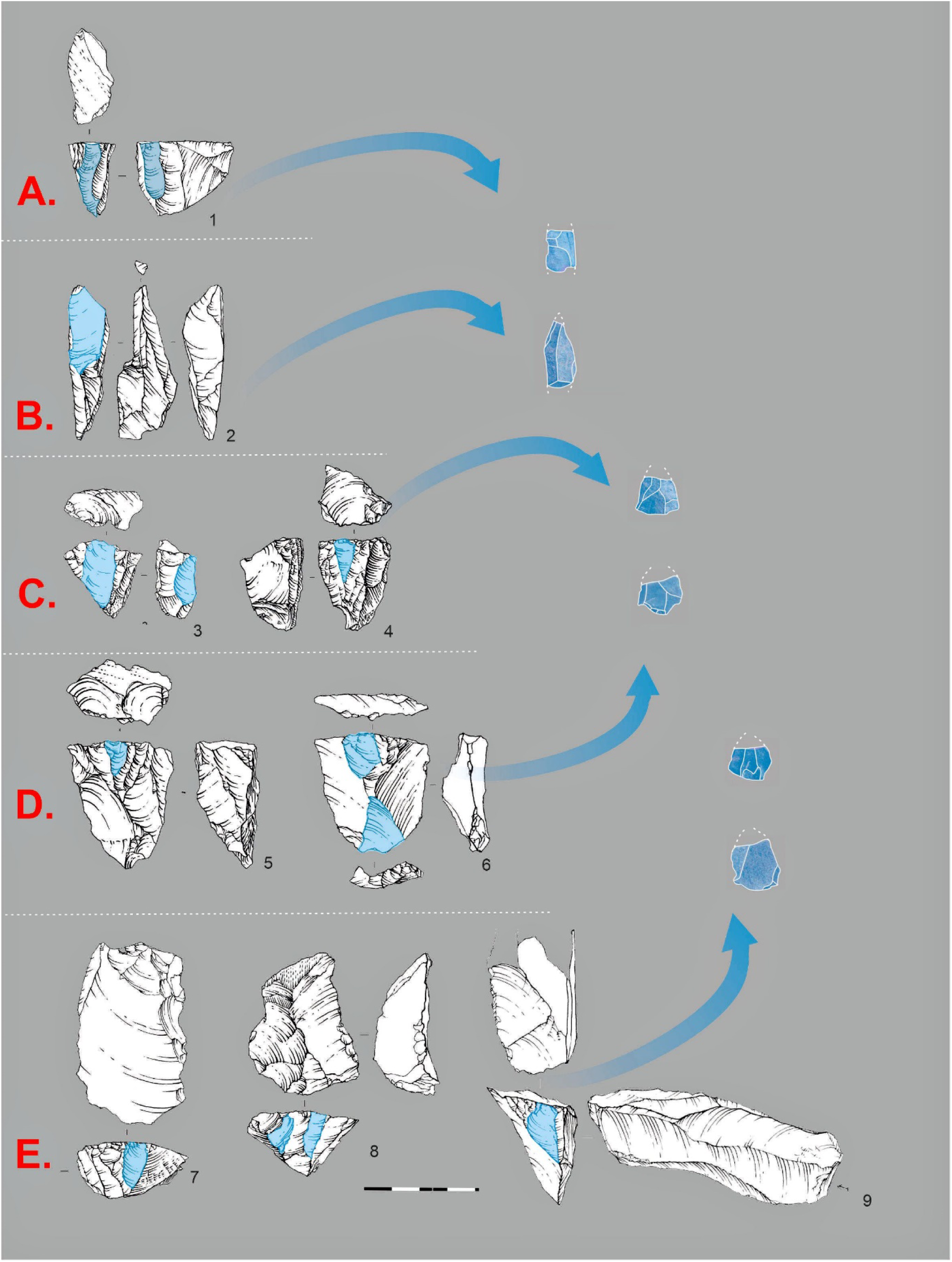
Technological chains of micro-point and bladelet production. A. 1 Narrow-faced core B. 2 Burin-cores. C. 3-4 Sub-prismatic, unidirectional cores. D Flat faced cores: 5 Flat faced, unidirectional; 6 Flat faced, bidirectional. E. 7-9 Carenated cores.

In addition, truncated faceted pieces (Fig 13) can be regarded as micro point cores. These pieces are notably present within the Obi-Rakhmat assemblage and were previously evaluated as tools in the context of layers 21-19 [47]. In other Middle Paleolithic complexes with evidence of the use of projectile armatures, analogous truncated-faceted pieces are considered as specialized cores for micro points [83,84]. We have conducted a comparative analysis of the angles between the flaking surface and the striking platform, using a range of specimen types, including blade cores, bladelet cores, and truncated faceted pieces. This analysis reveals that the angle of the truncated faceted pieces is notably sharper compared to that of the other cores (Fig 14). Furthermore, the dorsal angle of archaeological points where the proximal part is preserved demonstrates a greater alignment with the blade and bladelet core’s angle than truncated-faceted pieces (Fig 14). Therefore, it is hypothesized that some micro points may have been obtained from truncated-faceted pieces; however, it is rejected that such artefacts were systematically used as cores.

**Fig 13.**
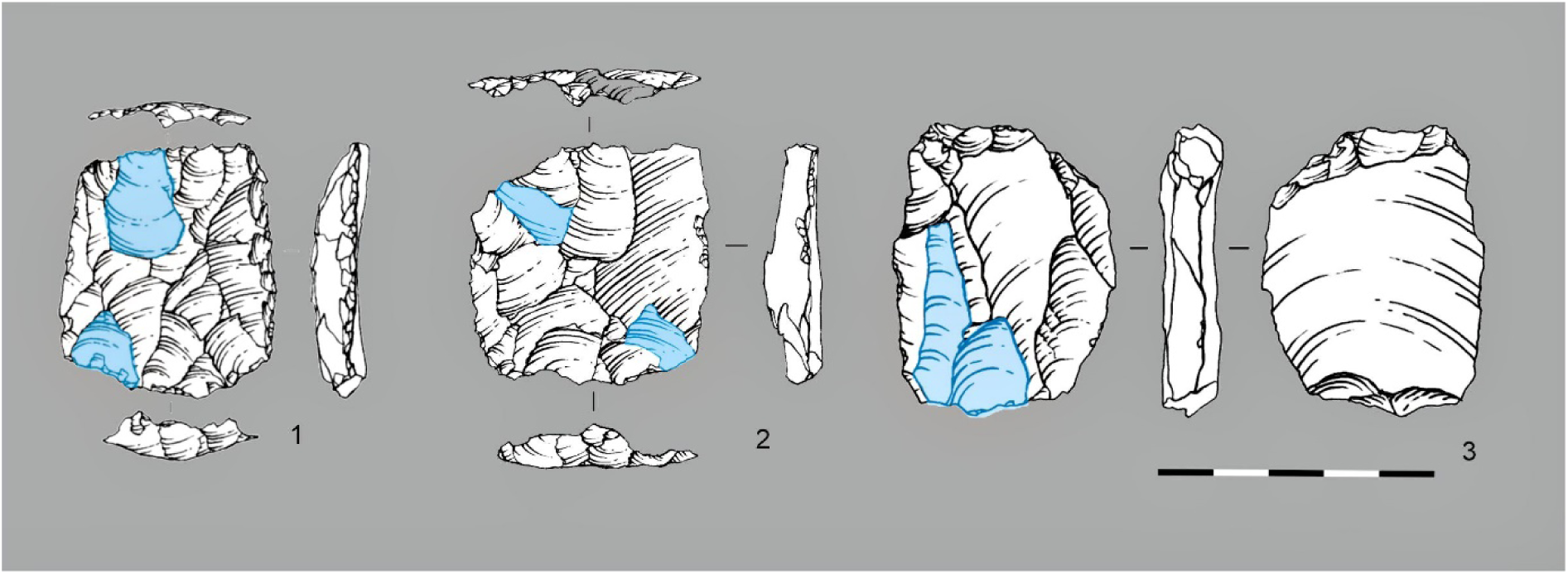
Truncated faceted pieces from layers 20-21: 1-3.

**Fig 14.**
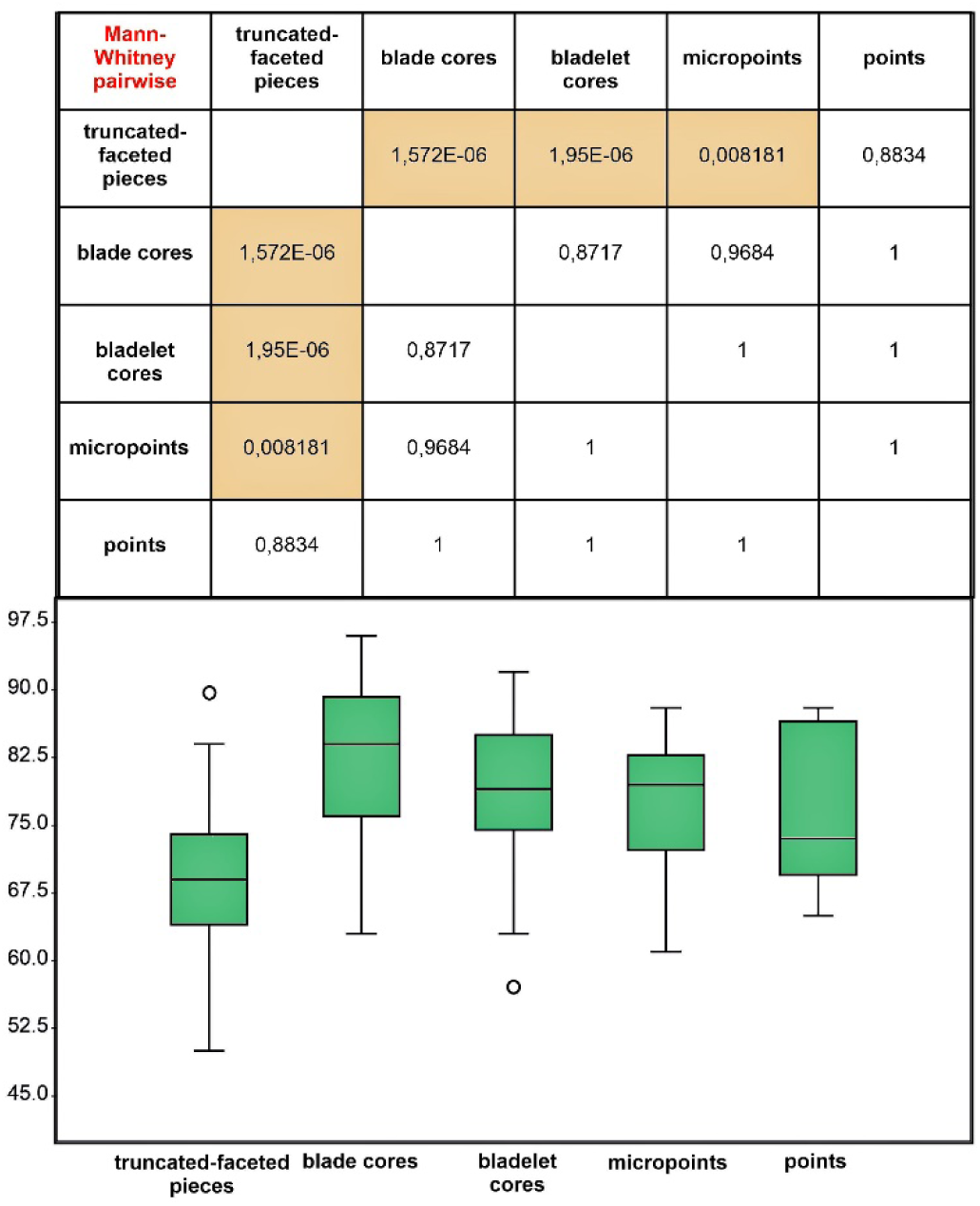
Comparative analysis of the angles between the flaking surface and the striking platform of cores, as well as the dorsal angle of points.

## Discussion

### Weapon identification

The question that usually arises with armatures is the type of weapon on which they are fitted. There are 6 different types of piercing weapon: dagger, hand-held stakes, hand-thrown spears, spears and darts thrown with a spear-thrower, arrows shot with a bow and blowpipe-fired darts. Apart from the dagger, which is an ancillary weapon for hunting, this classification according to distance of use also corresponds to the weight (and robustness) of the weapons related to the kinetic energy involved in their use, from the heaviest to the lightest. At the two ends of this range, the critical parameters are reversed: on the one hand, robustness is essential, while penetration is ensured in any case by the high power developed; on the other hand, the low impact power makes the sharpness of the projectile tip a determining factor. The design of the armatures will therefore depend on these constraints, in terms of shape and size. Numerous studies have attempted to find discriminating traceological criteria for distinguishing throwing techniques on the basis of the lithic points damage. They converge in concluding that the degree of damage is proportional to the energy involved [80,85,86], i.e. javelin points are more broken than arrowheads (all other things being equal), but none has set a threshold because there are too many parameters involved. Even admitting that the experimental conditions could be rigorously similar (weapon characteristics, type of hafting, method of throwing, shooting distance, hardness of the surrounding ground, ambient temperature, game, etc.) to the prehistoric ones - which is an illusion – a quantitative model would only be discriminating in a homogeneous archaeological assemblage in terms of all the parameters considered. The accumulation of points of the same type but fitted to both arrows and darts, or shot in summer and winter [87], is likely to make it inapplicable.

The only discriminating qualitative criterion, on ambivalent points (e.g. shouldered points), between arrows and spear heads, is the fracture by lateral flexion (from one edge to the other, not transversally) induced by the lever arm of a long shaft [4].

Recently, a new set of criterions has been highlighted, based on the ratio between the bending and compressive components of the point fracture according to the ballistic trajectory of the projectile. For the moment, however, the distinction is only relevant between hand-thrown javelins on the one side and arrows or darts on the other [73,88].

In any case, the current series from Obi-Rakhmat is much too small to apply any statistical model on traceological criteria. In order to hypothesize the type of weapon on which the points were mounted, we need to consider the technological parameters of their functioning. A 2 cm wide point weighing between 1 or 2 grams does not meet the same needs as a 4 cm wide point weighing 30 grams.

In the category of medium sized points, between tip crushed samples (Fig 5: 22, 28, S1 Fig) and a heavily laterally broken one (Fig5: 20), the range of energy involved is too wide for identifying a particular weapon, while their size could fit dart and spear. We can only note that the points which have the most cutting edges could have been mounted as daggers, a type of weapon that is rarely considered in the studies [79]. Unfortunately, the poor state of preservation does not allow for their detailed traceological analysis. Conversely, the narrow but thick point n°24 (Fig 5: 24), with less edge sharpness, seems to have been designed for a heavy dart or spear such as the broken one n°20 (Fig 5: 20).

The category of small points is more telling. Such small points were not designed to withstand violent impacts, their triangular shape made it impossible to bind them to the shafts, only to glue them, and these shafts could only be significantly smaller in diameter than their maximum cutting width [40]. It is important to remember that the role of the cutting armature is to tear the skin of the prey to open the way for the shaft, so that the skin does not tighten around it and reduce penetration. This is all the more important as the projectile has low momentum, which is the case for an arrow shot with a primitive bow. Contrary to what archaeologists have endlessly discussed, it is not the weight of the arrowhead that first matters but its penetration capacity, which is directly linked to its width and acuity for a given bow draw-weight [38,89–91].

Inventories of archaeological and ethnographic arrows, and even contemporary leisure and sporting archery, show that their shafts deviate very little from a diameter of 7-8 mm [92–96], depending on the power of the bow. Contrary to what Klaric et al. claim [97], this is not a “rule”, i.e. a cultural arbitrary, but the consequence of physical principles, i.e. a transcultural invariance. This is what A. Leroi-Gourhan called “trends”, as opposed to the “degrees of fact” [98].

This has to do with the flexibility (*spine*) of the arrow [73 Supplementary file2 video], which, if too great, will absorb part of the energy at launch and at penetration and induce too much oscillation during flight and at impact, or, if too weak, will not get around the bow handle and will deviate laterally, with, in both cases, an alteration in shot accuracy [38,40,99]. The dart’s spine [73 Supplementary file3 video,100] must be calibrated just as precisely, but according to the spear-thrower operating principle to which it contributes, with the same consequences for shot efficiency. One among the parameters used to adjust it is the resistance to acceleration induced by the mass of the tip [40,101].

Last but not least, resistance at penetration increases with the shaft’s diameter due to the greater surface area exposed to the tissues which results in a higher drag factor [38].

The difference in the principle of energy delivery between the bow and the spear-thrower, in an inverse relationship between velocity and mass, results in projectiles whose characteristics are not interchangeable and whose impact force are not equal (higher with darts). As written by Hughes “*Spine and size conformance (matching) between the projectile and the propulsive device are critical to insure accuracy and an efficient transfer of energy* [102]” [40].

This explains the bi-modal distribution distinguishing darts and arrows, according to head width and shaft diameter, reported by Marsh et al. [95 Fig. 4] from the measurements of 168 hafted points or shafts over whole American continent. The chosen geographical area, with its rich archaeological and ethnographic corpus, also offers the advantage of having known only one type of bow before the Hispanic period - the simple or self-bow - and the widespread use of composite arrows and darts made from a main shaft and a removable foreshaft. This technical solution is not exclusive to the American continent, as evidenced by the complete arrows from Egyptian Predynastic and Dynastic tombs (V and IVth millennium BC) whose lithic and bone armatures have typological equivalents in the Epipaleolithic and even the Late Stone Age [103]. The foreshaft reduces the dimensional gap between arrow and dart heads, by allowing to fix smaller points on composite darts than on single-piece ones. However, the dart foreshaft has to be more robust for enduring à greater stress than the arrow (greater impact force with more bending component [73]) which means a larger diameter.

In order to cut the skin widely enough for lowering the resistance to shaft penetration, Hughes calculates that the maximum circumference of arrow and dart tips should be 140-150% of the shaft diameter [40]. Applied to the complete micropoint (Fig 4: 00, Fig 8: 00, S17 Fig), the formula gives a value of 7 mm for its potential shaft, i.e. that of an arrow. Because of their quasi-equilateral outline, the triangular micropoints of Obi-Rakhmat have geometrically a lower cutting efficiency (*mechanical advantage*) than the Paleoindian points of thin elliptical cross section with much longer cutting edges from which this index has been extrapolated [40,104]. As a result, the corresponding theoretical shaft diameter should be seen as a maximum.

To refute the presence of arrowheads in contexts too ancient for considering the existence of bow- arrow technology, Klaric et al. have recently proposed the alternative hypothesis of reduced weapons for children with which the criterion of efficiency becomes secondary [97]. On the basis of ethnographic data from three continents, they distinguish two cases that can be archaeologically detected: simplified lithic or bone points expeditiously made by adults or clumsy ones made by the children themselves. It’s true that the place of children in prehistoric societies is not considered a research topic in itself, even though its anthropological implications would be fundamental, however the proposed criteria don’t fit in with Obi-Rakhmat’s industry. The micropoints there are not reduced replicas of larger points resulting from a same debitage scheme (Fig 15) and their experimental reproduction requires a real know-how. To presume that children were learning lithic knapping by using their own solutions, different from those practiced by adults, is not the kind of parsimonious hypothesis advocated by Klaric et al. For the time being, the most simple hypothesis is to consider the production scheme of the micropoints at Obi-Rakhmat as a response to the specific need for small, light-weight sharp armatures.

**Fig 15.**
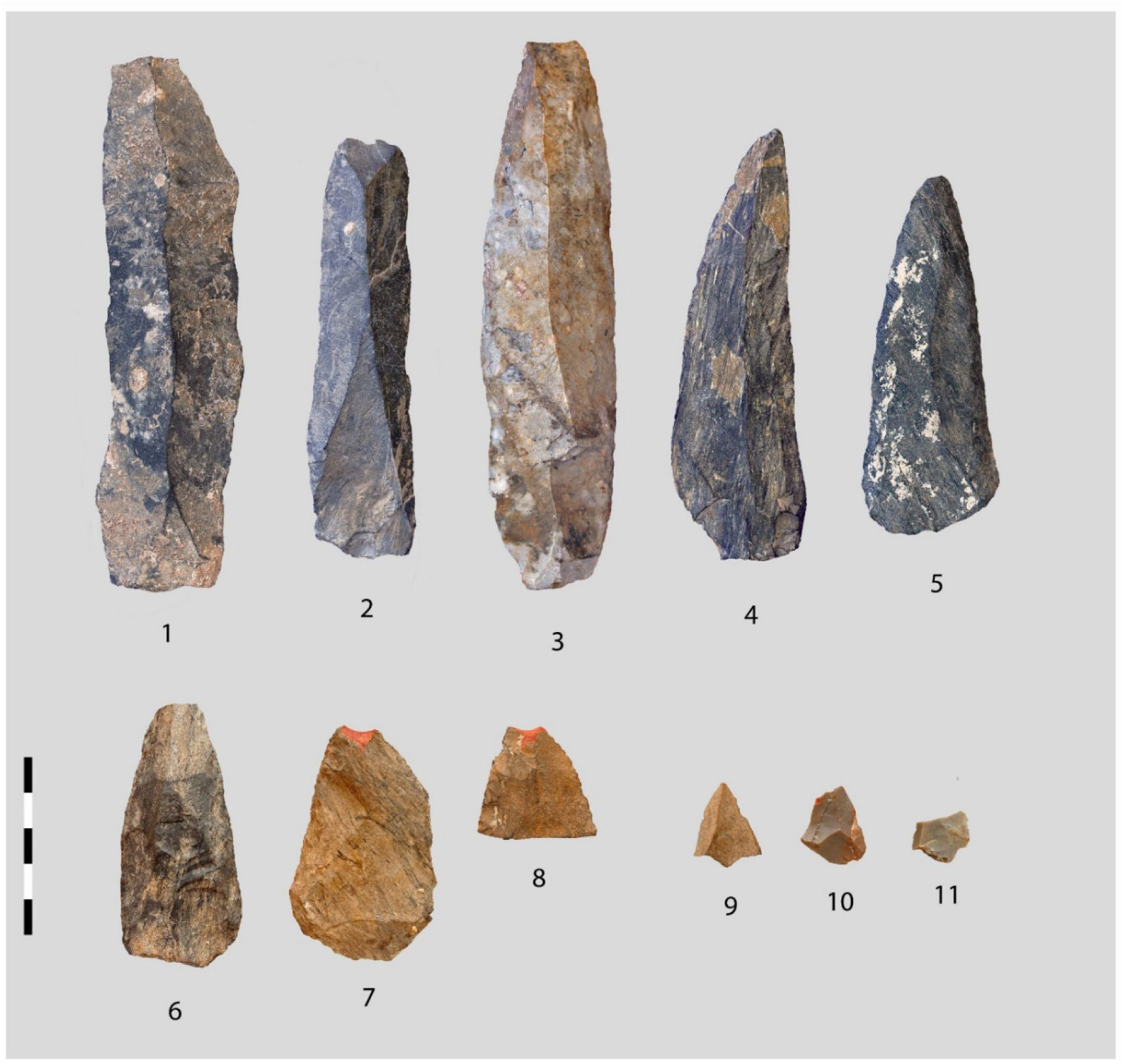
**Micropoints alongside the structuring products of the lithic industry**. 1-5 blades and elongated points from blade cores, 6-8 Medium sized retouched weapons heads, 9-11 Micropoints. Centimetric scale.

A technical alternative for such small projectile points, but which to our knowledge has not been experimentally tested, could be that of a dart armature shot with a blowgun. However, the ethnographic examples documented in South-East Asia and in the American continent [105], originated from a core area in tropical forest, do not mention darts reinforced with lithic or bone point. The only documented composite darts are Malayi ones from South India, with a detachable conical barbed steel head, linked to a short shaft by a long and thin cord wound closely around it, which were used for shooting fishes at short distance [106]. The Cherokee blowgun can deliver a dart made from a very light wood or bamboo splinter at nearly twice the velocity of an arrow from a native self-bow [107]. Its effective hunting range is between 12 and 18 m [108,109]. The darts of Jahai people in Malaysia, among the most ancient blowgun hunters, weight less than one gram and can be shot precisely at 50 m [110]. On such needle-like projectiles, micropoints from Obi-Rakhmat would be in every respect (width, fitting, weight distribution, inertia, ballistic properties) out of proportion.

### Comparison with former, contemporary or similar projectile heads

The earliest lithic weapon armatures reported in the literature on a traceological basis are those from Kathu Pan 1 in South Africa, stratum 4a, which has yielded the oldest dated Fauresmith Industry [111–114], at the transition between Earlier and Middle Stone Age, ca 500 ka. This KP1 assemblage contains numerous unifacially retouched points and nonretouched triangular flakes and blades from Levallois knapping. In a sample of 210 points and fragments covering 4 square units, 31 *diagnostic impact fractures* (DIF) were detected on 29. However, only 5 macroscopic photos of these DIF, including 2 tiny burin removals, are provided but no view of the whole artefacts (photo nor drawing). These possible spearheads are not said to differ in size or morphology from the rest of the corpus. The average length of all the points (retouched or unretouched, regular or not) is over 7 cm. No human remains have been found with this ancient expression of Fauresmith-type industry.

A little less ancient (> 279 ka) are the obsidian points from the Gademotta site complex in Ethiopia, identified as projectile points from velocity-dependent microfracture features, diagnostic damage patterns, and artifact shape [34], however, the interpretation has been controversial, with the apical removals being seen as resulting from a sharpening process by lateral tranchet blow removals rather than from projectile impact [35,115].

The lithic industry of the Misliya cave terrace, in Israel, associated with the oldest remains of archaic Sapiens found outside Africa (ca. 180 ka), consists mainly of points of various types (unretouched Levallois, retouched Levallois and Abu-Sif and Misliya points). Three traceological analyses were carried out on samples of the lithic assemblage containing the different types of points, i.e. a total of 344 points out of a global sample of 445 pieces. The aim of the first study was to find weapon heads [72], while the next 2 were exhaustive [46,116]. In all, 48 weapon heads were identified on the basis of their microscopic damage and microscopic linear impact traces (5% of the sample). The dominant type are the unretouched Levallois points with 26 specimens. Their average length is 64 mm and their average weight is 19 grams. Nevertheless, the complete study shows that the points are at Misliya multifunctional tools also related to the acquisition of vegetal foods as well as performing craft-related activities.

Only slightly more recent are the 3 non-tanged projectile points identified in the lower section of the stratigraphy of the Ifri n’Ammar rock shelter in Morocco, dated between 143 and 171 ka [117]. They precede the tanged Aterian points of the upper section, dated between 171 to 83 ka, of which 11 out of 42 have also been identified as spear tip on the basis of their damage [118]. The way they were hafted and their size do not seem to distinguish them from common tools. They are more massive and robust than the micropoints of Obi-Rakhmat.

On the other side of the Mediterranean, further north, in the world of the Neanderthals, at the Bouheben site in France, in an assemblage of 125 Mousterian points from around the same time period (geological assignment to MIS 6) as Misliya, 6 specimens with impact scars interpreted as weapons head are mentioned. [119]. Four fairly convincing macro photographs are published, but no views of the whole specimens. This suggests that there was nothing morphologically distinctive that was worth showing.

In the rich Levallois Levallois industry from level IIA of Biache-Saint-Vaast, northern France, a little older (MIS 7), 16 spear points and 20 butchering knives have been identified among the Mousterian points and convergent scrapers by a macroscopic and microscopic traceological analysis of 157 convergent pieces, side scrapers and Levallois implements [120]. The weapons heads are among the most elongated and symmetrical samples in the corpus.

The macroscopic analysis of 119 nonretouched Levallois points from the slightly younger assemblage of Therdonne (France, MIS 7/6) has been less successful, providing only 2 possible weapons heads but 17 butchering knives [121,122].

Although a hundred thousand years more recent, the assemblages from Angé and Bettencourt-Saint-Ouen, in the northern half of France, do not present a different pattern, with triangular-shaped pieces meeting a variety of needs, among which weapon point is just one use. At Angé, lithic points were mainly produced by a convergent unipolar production scheme, but also by bipolar and centripetal Levallois schemes [123]. One weapon point has been noticed because of its lateral damage [124], however macroscopic and microscopic analysis of 33 other Mousterian points of different sizes and morphologies revealed only harvesting and plant-working tools (Plisson, unpublished). At Bettencourt-Saint-Ouen, where points were also produced by different schemes, the analysis of 49 Levallois points has uncovered 1 weapon point with lateral scars, 8 butchering knives and 2 wood knives [Caspar in 125]. These two sites are contemporary with Obi-Rakhmat’s level 21, but have nothing in common with its micro projectile points.

To find small projectiles heads, it is necessary to leave Neanderthals, cross back the Mediterranean Sea and go as far as the southern end of Africa. Between > 77ka and 64ka, across 3 distinct cultural assemblages, the Sibudu rock shelter, in the province of KwaZulu-Natal, displays a range of types and sizes of projectile points, from Pre Still Bay serrated bifacial lithic ones to Howiesons Ports microlith quartz segments and bone points, part of which being described as arrowheads [119,126–138]. Lithic segments and bone points continued to be used locally as arrowheads well into historic times [139–141].

However, the very different lithic head designs (Levallois and pseudo Levallois *vs* serrated bifacial, foliated or segments), the dates, the distance and the absence of geographical relays prevent from seeing any relationship between Obi-Rakhmat earliest levels and the different cultural layers of Sibudu. The contribution of South Africa to the expansion of Anatomically Modern Human (AMH) is more likely to have been in the direction of Asia, according to the coastally oriented dispersal model, due, among other arguments, to the similarity of the Howiesons-type segments with those of the first microlithic assemblages from India and Sri Lanka [142].

The closest technical comparison is not in Africa, but in the Rhône valley in France, in a brief Neronian occupation of the Mandrin cave 54,000 years ago (51.7-56.8 interval). This cultural layer E yielded micropoints identical to those found at Obi-Rakhmat, also impacted by use as projectile heads (Fig 16). Due to their tiny size, they are interpreted as arrowheads [7]. A Homo Sapiens deciduous tooth has been found in the same layer [143]. A later local Mousterian layer, with Neanderthal mandibular remains, closes the Middle Palaeolithic stratigraphic sequence at the site [144]. The Neronian with its small arrowheads is regarded as belonging to a first pioneering wave of AMH incursion into southern Europe [145]. Other sites with lithic industries using the same reduction strategy and characterized by the combination of axial-macro and micropoints/blades/bladelets are beginning to be documented at the end of the Middle Palaeolithic in Spain with the Arlanzian of Cueva Millán [146] and in Italy at Riparo l’Oscurusciuto [147]. At Cueva Millán micropoints impacted by their potential use as projectile head are mentioned, while a past study at Riparo l’Oscurusciuto highlighted the presence of impact scars on 6 macro-points, suggesting their use as tip spears [148]. The biological identity of these innovative hunters is unknown, but so far, the combination of different sizes of projectile heads and the use of microlithic ones is not part of the Neanderthal repertoire [149], while the resemblance with the Neronian suggests that the same author could be involved.

**Fig 16.**
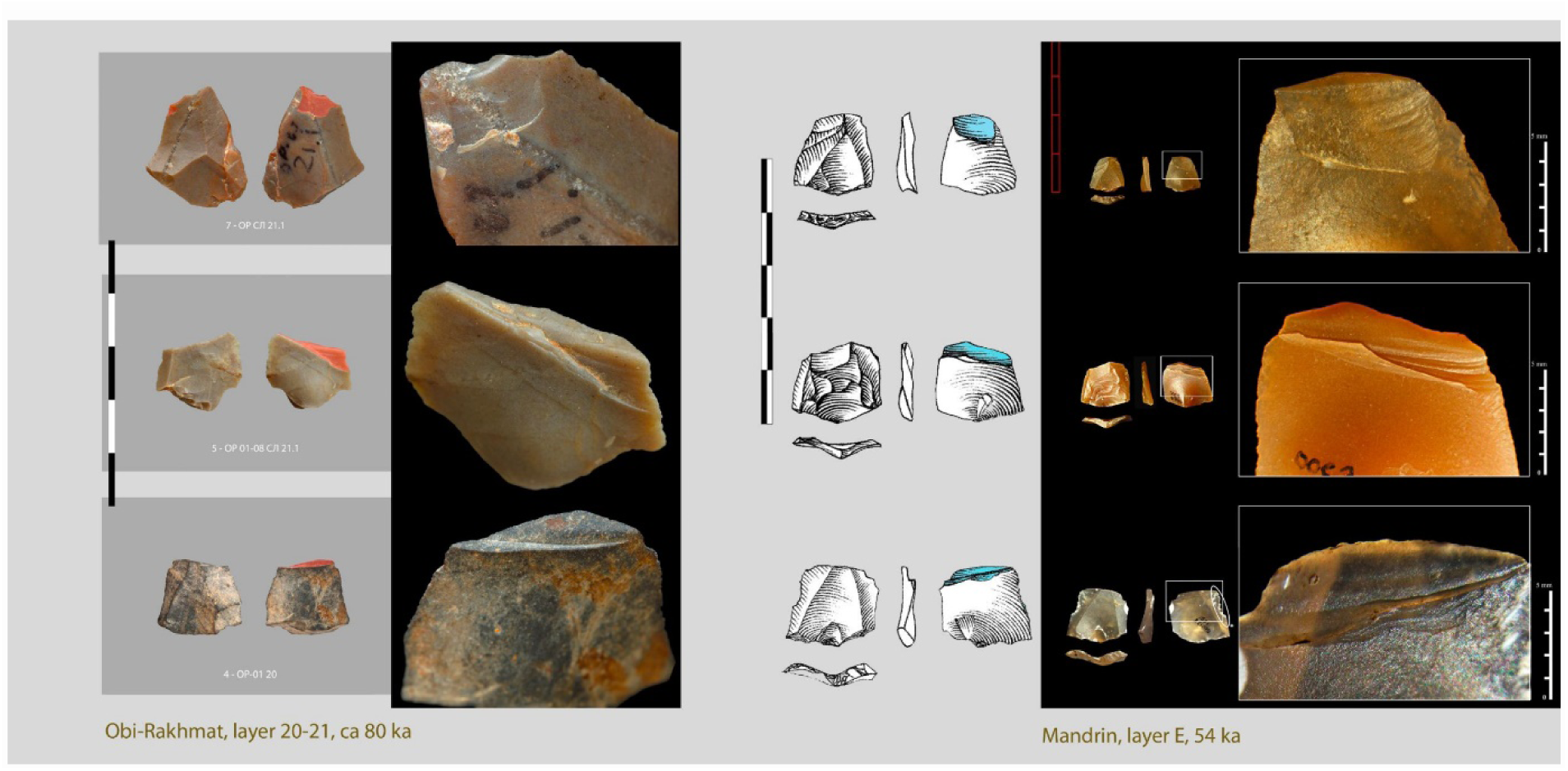
Comparison between micropoints from Obi-Rakhmat, layers 20-21, and Mandrin, layer E [83,150].

### The IUP perspective

A parallel is drawn between the industries of layer E at Mandrin and layer XXIV at Ksar Akil in Lebanon, which share notable technical similarities [143]. Due to the early excavation conditions, smallest micropoints are lacking at Ksar Akil, while, according to the dating, there is a gap of almost 10,000 years between the two assemblages. This gap is marked in the stratigraphy of Ksar Akil by an abrupt break between layers XXVI and XXIV, without possibilities of technological continuity between the Middle Palaeolithic and the Initial Upper Palaeolithic [145] and by a partial maxilla of young AMH [151]. The absence of AMH specimen in the Levant since the material from Skhul/Qafzeh (∼120–90 ka ago), except the calvaria from Manot Cave in Galilee dated to 54.7 ± 5.5 ka [152,but see 153], does not advocate for a continuous representation and local evolution of AMHs in the Levant. The fact that the earliest dates for the IUP layers of Ksar Akil post-date the Neanderthal era could suggest that earlier settlements of AMH returning to the Levant did not take place on sites that were at the time occupied by Neanderthals. More generally, we might ask whether recent developments in dating and calibration techniques are making it possible to compare, in absolute terms, results spread over almost a decade in different contexts?

Whatever the moment of return to the Levant of the AMHs, considered to be the artisans of the IUP, the milestones on the evolution of hunting weapons that would make it possible to draw the link are for the moment lacking in East Africa or Arabia. In the Levant itself, the projectile head design known from Um El Tlel or Nahal at around 60 ka is no different from the rest of the toolkit like in any former Neanderthal site [23,79,149,154–156]. That’s why Ludovic Slimak suggests to look further east for the possible roots of the very first pioneer groups of Sapiens to enter Europe [145].

Despite the distance in time (∼ 25 ka) and space (∼ 6000 km) that separates the first occupations of Obi-Rakhmat from the layer E of Mandrin, we can only be surprised by the great morphological similarity of their micro-projectile points, which look interchangeable. As well as at Cueva Millán and Riparo l’Oscurusciuto, they are resulting from comparable specific flaking patterns, which are characterised by their flexibility. The massive blades of Obi-Rakhmat are absent from the more recent assemblages, but all four are characterised by Upper Palaeolithic-type laminar volumetric cores, and by the production of axial points, blades and bladelets.

We do not pretend to draw a direct link between Obi-Rakhmat and Mandrin, but these astonishing similarities, particularly in one element relating to a complex weapon system, of which this could be one of the earliest expressions, give rise to a number of reflections. In the same way that the technical solutions developed around 64 ka at Sibudu, in South Africa, lasted well into the Holocene, well beyond the particular cultural group that invented them, it is likely that micropoints of the type described here and what they encompass spread between different groups, as bow and arrows are particularly well suited to hunting game that is difficult to approach in an open foothill environment, such as the Asian ibex, and to broadening the spectrum of hunted fauna.

On the basis of current genetic data the Persian plateau, at the north-eastern periphery of which Obi-Rakhmat is located, has been recently defined as a population hub where the ancestors of all present-day non Africans lived between the early phases of the Out of Africa expansion (∼70–60 ka) and the broader colonisation of Eurasia (∼45 ka) [157]. This resource-rich environment, with its diversity of topography and water sources [158], provided a refuge area conducive to demographic regeneration after the Out of Africa bottleneck, and consequently to interaction between groups. These interactions probably encouraged technical innovations in response to the new environmental variables, including projectile weapons [157].

The visibility of the IUP in the Levant and Europe from ∼ 45 ka may be the consequence of a same population spreading from the Persian plateau [the second wave in the model proposed by 145]. This could explain the morphological difference between the light projectile armatures of the Emiran/IUP sequence from Boker Tachtit [9] and those from Mandrin or Obi-Rakhmat, which would represent an initial, i.e. more archaic, phase. This is suggested by the fact that the shape of the micropoint was obtained in various ways. It was not yet the primary objective around which to organize the productions schemes. Light projectile points, whose functional design impose standardisation, were initially added on the margin of pre-existing production systems centred on other requirements. This corresponds to what the philosopher Gilbert Simondon [159,160] sees as the birth of a new technical lineage, whose initial stages of concretising the new principle on which it is based borrow from the technical environment in which it emerges, before tending towards the conditions of its own coherence. By becoming the structuring element in tool production, due to their importance in the subsistence system, projectile points, through their specialization and standardization, will probably influence Sapiens’ technical conceptions in a way that will become his signature.

At this point, the question that can no longer be avoided is that of the biological identity of Obi-Rakhmat’s inhabitants. In 2003, teeth and skull fragments from a single juvenile individual (9-12 years old) were discovered in layer 16 (ca 70 ka) which express a relatively Neandertal-like dentition but coupled with more ambiguous cranial anatomy [48,65,161–163]. The most immediate assumption would be that it was a Neanderthal-[163] or Denisovian-Sapiens hybrid, which could explain its premature death.

The only known association of Neanderthal remains with different types of weapon points, in this instance lithic and bone points, is in the Castelperronian of Arcy sur Cure cave in France. However, contamination with the underlying Mousterian level, which was rich in Neanderthal remains, and the recent identification of an anatomically modern ilium of a neonate [164], call such an association into question [165], while at Saint Cézaire, in France, the combination of Castelperronian lithic industry and human remains seems to be the result of solifluction [166]. As for knowledge of Denisovian technical systems, it is currently based on a single site where the only projectile element identified to date is synchronous with the presence of AMH [167,168].

If we consider the diversification of projectile armatures and the invention of propulsion instruments with their inherent complexity (systemic integration of a large number of element) [169] as markers of a development specific to Sapiens, we need to reformulate the question of the transition between the Middle Palaeolithic and the Upper Palaeolithic, not in biological terms, between hominins (in fact, historiographically between Neanderthals and Sapiens, the Denisovians being excluded despite their genetic legacies) but in relation to the steps in the trajectory of AMH, probably of a demographic nature [170]. This would give back to the word *transition* the notion of continuity that it conveys.

For now, the absence of genomic or proteomic analysis at Obi-Rakhmat leave the question open. Whatever the reply, it will be of the utmost importance to the debate on the roots of the IUP.

## Conclusion

The results presented here are based on a preliminary study on material from the oldest layers of Obi-Rakhmat, which is currently being collectively studied. The number of pieces identified as projectile points may seem low, but it only covers 20 square metres and, according to an agent-based modelling, could be in fact regarded as high [171].

In any case, the sample is sufficient to highlight the presence of micropoints impacted by use as projectile heads (including two with a combination of macroscopic and microscopic traces) in these layers dated to ∼ 80,000 years ago. Their almost microlithic shape makes them technically unsuitable for mounting on anything other than arrow shafts. The same type of armature is described in a pioneer settlement by Sapiens in the Rhône Valley 25,000 years later.

At Obi-Rakhmat, they are present in the inventory with more robust points and with bladelets. These different elements seem hard to match on the same shaft and probably corresponded to 3 types of weapons.

The bladelets will need a dedicated functional study to see if they were commonly used as projectile barbs. The next task will be to see whether the same technical solutions are present throughout the stratigraphy or whether this combination of weapons was specific to the first layers of occupation. The question is especially relevant for the micropoints, which are about to become an index fossil. So far unnoticed because they are unretouched, tiny and fragmentary, it is likely that they will now start to appear in sites between Central Asia and Western Mediterranean Sea. If so, it will be meaningful to see how Obi-Rakhmat fits into the dynamics of their dissemination: as a recipient site or as part of the cultural complex in which they were invented? The micropoints could be an indicator of interconnections between distinct groups and of their temporality in relation to climatic phases.

The difficulty of integrating typological, technological, chronological, geographical and anthropological data relating to the Initial Upper Palaeolithic and its premices into an unifying model suggests that what this concept encompass, linked to the expansion of Sapiens into Eurasia, was not a sudden linear phenomenon but the result of a mosaic of interactions over millennia between groups emerging from Africa and vernacular populations [172]. Small projectile points and the complex weaponry behind can provide a discriminating criterion in time and space for getting a clearer view.

## Supporting information

The photographic plates include a double-sided view of the artefacts, with features considered diagnostic coloured red, as well as macroscopic views of these features.

Due to the file size limit, only the 3D models of most relevant artefacts are included in this draft. Their content is activated by clicking.

S1 Table. Technological inventory from layers 20 and 21 (PDF).

S1 Fig. Retouched point 26 - ОР 10 06 21.1 622, photographic view, centimetric scale. (JPG).

S1 Fig. Retouched point 26 - ОР 10 06 21.1 622, 3D model (PDF).

S2 Fig. Retouched point 28 - ОР 11 21.2 06 КВ 145, photographic view, centimetric scale (JPG).

S2 File. Retouched point 28 - ОР 11 21.2 06 КВ 145, 3D model (PDF).

S3 Fig. Apical fragment of retouched point 22 - ОР 11 21.1, photographic view, centimetric scale (JPG).

S3 File. Apical fragment of retouched point 22 - ОР 11 21.1, 3D model (PDF).

S4 Fig. Apical fragment of retouched point 104 - OP 01 19 4, photographic view, centimetric scale (JPG).

S4 File. Apical fragment of retouched point 104 - OP 01 19 4, 3D model (PDF).

S5 Fig. Retouched point 24 - ОР СЛ 21 1 196, photographic view, centimetric scale (JPG).

S5 File. Retouched point 24 - ОР СЛ 21 1 196, 3D model (PDF).

S6 Fig. Broken retouched point 20 - ОР 10 21.1 0-6, photographic view, centimetric scale (JPG).

S6 File. Broken retouched point 20 - ОР 10 21.1 0-6, 3D model (PDF).

S7 Fig. Apical fragment of retouched point 30 - ОР 11 21.2 П6 17, photographic view, centimetric scale (JPG).

S7 File. Apical fragment of retouched point 30 - ОР 11 21.2 П6 17, 3D model (PDF).

S8 Fig. Apical fragment of retouched point 27 - ОР 11 П 7 21.1, photographic view, centimetric scale (JPG).

S8 File. Apical fragment of retouched point 27 - ОР 11 П 7 21.1, 3D model (PDF).

S9 Fig. Broken micropoint 4 - OP-01 20, photographic view, centimetric scale (JPG).

S9 File. Broken micropoint 4 - OP-01 20, 3D model (PDF).

S10 Fig. Broken micropoint 5 - ОР 01-08 СЛ 21 1, centimetric scale (JPG).

S10 File. Broken micropoint 5 - ОР 01-08 СЛ 21 1, 3D model (PDF).

S11 Fig. Broken micropoint 7 - ОР СЛ 21.1, photographic view, centimetric scale (JPG).

S11 File. Broken micropoint 7 - ОР СЛ 21.1, 3D model (PDF).

S12 Fig. Broken micropoint 8 - ОР 11 П 7 21, photographic view, centimetric scale (JPG).

S12 File. Broken micropoint 8 - ОР 11 П 7 21, 3D model (PDF).

S13 Fig. Broken micropoint 9 - ОР 01 СЛ20.3, photographic view, centimetric scale (JPG).

S13 File. Broken micropoint 9 - ОР 01 СЛ20.3, 3D model (PDF).

S14 Fig. Broken micropoint 11 - ОР 2001-2008 СЛ 21.1, photographic view, centimetric scale (JPG).

S14 File. Broken micropoint 11 - ОР 2001-2008 СЛ 21.1, 3D model (PDF).

S15 Fig. Broken micropoint 16 - ОР 2001-2008 СЛ 21.1, photographic view, centimetric scale (JPG).

S15 File. Broken micropoint 16 - ОР 2001-2008 СЛ 21.1, 3D model (PDF).

S16 Fig. Broken retouched micropoint 2 - ОР 21.1 280, photographic view, centimetric scale (JPG).

S16 File. Broken retouched micropoint 2 - ОР 21.1 280, 3D model (PDF).

S17 Fig. Levallois micropoint 00 - OP Сл 21 X7, photographic view, centimetric scale (JPG).

S17 File. Levallois micropoint 00 - OP Сл 21 X7, 3D model (PDF).

S18 Fig. Broken retouched bladelet 19 - ОР 11 СЛ 21.1 П8, photographic view, centimetric scale (JPG).

S18 File. Broken retouched bladelet 19 - ОР 11 СЛ 21.1 П8, 3D model (PDF).

S19 Fig. Broken burnt crushed bladelet 15 - ОР 11 21.1 07, photographic view, centimetric scale (JPG).

S19 File. Broken burnt crushed bladelet 15 - ОР 11 21.1 07, 3D model (PDF).

S20 Fig. Backed bladelet 31 - ОР-08 Сл 21.1 КВ М7, photographic view, centimetric scale (JPG).

S20 File. Backed bladelet 31 - ОР-08 Сл 21.1 КВ М7, 3D model (PDF).

S21 File. Broken micropoint 6 - ОР 2001-2008 СЛ 21.1, 3D model (PDF).

S21 Fig. Experiments by Vladimir Kharevich, photographic view (JPG).

## Supporting information

Supplemental Illustrations

## Acknowledgements

We are grateful to Dr. Farhod A. Maksudov, Director of the National Archaeological Center, Academy of Sciences of the Republic of Uzbekistan, and all the staff at the for facilitating access to the collection.

Special thanks to Dr. Françoise Courmelon and Dr. Colline Bataille for making possible the participation of H.P. to this study.

## Contributions

**Conceptualization**: Hugues Plisson

**Data Curation:** Ksenya Kolobova, Alëna Kharevich

**Formal Analysis:** Ksenya Kolobova

**Funding Acquisition:** Andrei Krivodhapkin, Hugues Plisson

**Investigation:** Hugues Plisson, Valdimir Kharevich, Alëna Kharevich, Lydia Zotkina, Malvina Bauman, Andrei Krivodhapkin

**Methodology:** Hugues Plisson, Valdimir Kharevich

**Project Administration:** Andrei Krivodhapkin

**Resources:** Andrei Krivoshapkin, Ksenya Kolobova, Hugues Plisson

**Software:** Pavel Chistyakov

**Supervision:** Andrei Krivoshapkin

**Validation:** Andrei Krivoshapkin, Ksenya Kolobova, Hugues Plisson

**Visualization:** Hugues Plisson, Pavel Chistyakov, Lydia Zotkina, Malvina Bauman, Eric Pubert

**Writing – Original Draft Preparation:** Hugues Plisson, Alëna Kharevich

**Writing – Review & Editing:** Andrei Krivoshapkin, Malvina Baumann

