## Supplemental Illustrations for "Arrow heads at Obi-Rakhmat (Uzbekistan) 80 ka ago"

Supplementary Tables 1

### Obi-Rakhmat, layers 20-21

Technological inventory

| | 20 | | | 21,1 | | | 21,2 | | | 21,3 | | |
|---|---|---|---|---|---|---|---|---|---|---|---|---|
| | N | % | %* | N | % | %* | N | % | %* | N | % | %* |
| **Cores** | 14 | 0,42% | **1,64%** | 72 | 1,30% | **2,49%** | 9 | 0,46% | **1,41%** | 2 | 0,34% | **0,97%** |
| **Core-like** | 5 | 0,15% | **0,59%** | 43 | 0,77% | **1,48%** | 2 | 0,10% | **0,31%** | 6 | 1,01% | **2,91%** |
| **Technical flakes** | 21 | 0,6% | **2,5%** | 44 | 0,8% | **1,5%** | 17 | 0,9% | **2,7%** | 8 | 1,3% | **3,9%** |
| **Blades** | 280 | 8,5% | **32,8%** | 1265 | 22,8% | **43,7%** | 216 | 11,1% | **33,9%** | 64 | 10,7% | **31,1%** |
| **Bladelets** | 257 | 7,8% | **30,1%** | 613 | 11,0% | **21,2%** | 152 | 7,8% | **23,9%** | 44 | 7,4% | **21,4%** |
| **Microblades** | 56 | 1,7% | **6,6%** | 109 | 2,0% | **3,8%** | 9 | 0,5% | **1,4%** | | 0,0% | **0,0%** |
| **Points** | 12 | 0,4% | **1,4%** | 120 | 2,2% | **4,1%** | 43 | 2,2% | **6,8%** | 19 | 3,2% | **9,2%** |
| **Flakes > 3 cm** | 208 | 6,3% | **24,4%** | 630 | 11,4% | **21,8%** | 189 | 9,7% | **29,7%** | 63 | 10,6% | **30,6%** |
| ***Including Tools:*** | *32* | 1,0% | **3,8%** | *209* | 3,8% | **7,2%** | *34* | 1,7% | **5,3%** | *9* | 1,5% | **4,4%** |
| **Flakes 1–3 cm** | 589 | 17,9% | **-** | 1533 | 27,6% | **-** | 1116 | 57,1% | **-** | 304 | 50,9% | **-** |
| **Chunks/shatter** | 48 | 1,5% | **-** | 57 | 1,0% | **-** | 34 | 1,7% | **-** | 19 | 3,2% | **-** |
| **Chips** | 1806 | 54,8% | **-** | 1064 | 19,2% | **-** | 166 | 8,5% | **-** | 68 | 11,4% | **-** |
| | **3296** | **100,0%** | | **5550** | **100,0%** | | **1953** | **100,0%** | | **597** | **100,0%** | |

Table. Overview of the lithic assemblage

| **CORES** | **20** | | **21,1** | | **21,2** | | **21,3** | |
|---|---|---|---|---|---|---|---|---|
| | **N** | **%** | **N** | **%** | **N** | **%** | **N** | % |
| **Blade cores** | 2 | 14,3% | 24 | 33,8% | 2 | 22,2% | 1 | 50,0% |
| **Levallois (points cores)** | | | 4 | 5,6% | 3 | 33,3% | | |
| **Levallois (flakes cores)** | 1 | 7,1% | 5 | 7,0% | | | | |
| **Bladelet cores** | 7 | 50,0% | 25 | 35,2% | 3 | 33,3% | | |
| **Flake cores (other systems)** | 4 | 28,6% | 13 | 18,3% | 1 | 11,1% | 1 | 50,0% |
| **Total** | 14 | | 71 | 100,0% | 9 | 100,0% | 2 | 100,0% |

Table.  Type of determinable  cores (core-like pieces are excluded)

| CORES | 20 | | 21,1 | | 21,2 | | 21,3 | |
|---|---|---|---|---|---|---|---|---|
| | N | % | N | % | N | % | N | % |
| **Flat faced, Unidirectional** | - | - | 6 | 25% | - | - | 1 | 100% |
| **Flat faced, Bidirectional** | - | - | 3 | 13% | - | - | - | - |
| **Sub-prismatic Unidirectional** | - | - | - | - | 2 | 100% | - | - |
| **Sub-prismatic Bidirectional** | - | - | 1 | 4% | - | - | - | - |
| **Sub-prismatic Asymmetrical (*Semi-tourné*)** | - | - | 2 | 8% | - | - | - | - |
| **Narrow-faced** | 2 | 100% | 12 | 50% | - | - | - | - |
| **Total** | **2** | **100%** | **24** | **100%** | **2** | **100%** | **1** | **100%** |

Table. Type of Blade cores

| CORES | 20 | | 21,1 | | 21,2 | | 21,3 | |
|---|---|---|---|---|---|---|---|---|
| | N | % | N | % | N | % | N | % |
| **Carenated** | | | 4 | 16,0% | | | | |
| **Narrow-faced** | 1 | 14,3% | 4 | 16,0% | 2 | 66,7% | 0 | |
| **Burin-cores** | 3 | 42,9% | 5 | 20,0% | | | | |
| **Flat faced** | 1 | 14,3% | | | | | | |
| **Sub-prismatic** | | | 7 | 28,0% | | | | |
| **Other core on flakes** | 2 | 28,6% | 5 | 20,0% | 1 | 33,3% | | |
| **Total** | 7 | 100,0% | 25 | 100,0% | 3 | 100,0% | 0 | |

Table. Bladelet cores

S1 Fig

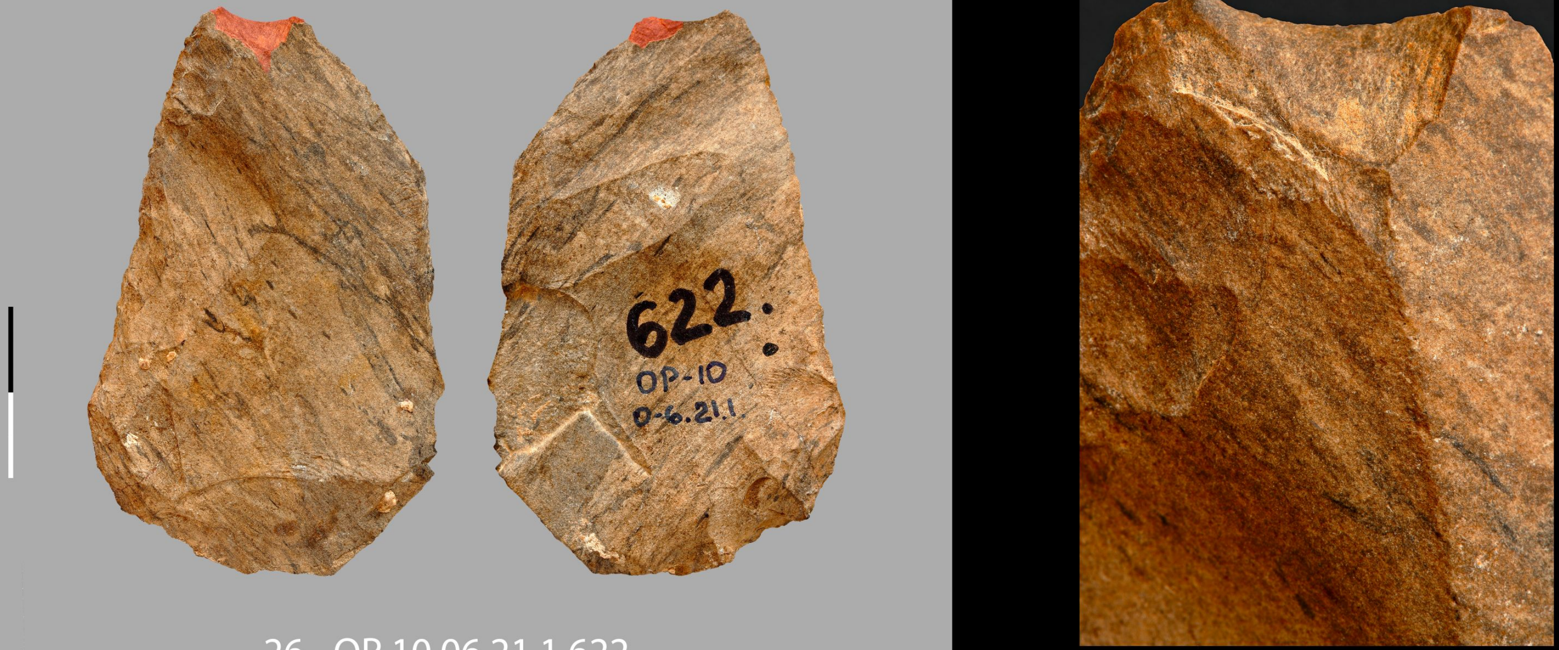


26 - OP 10 06 21.1 622

S2 Fig

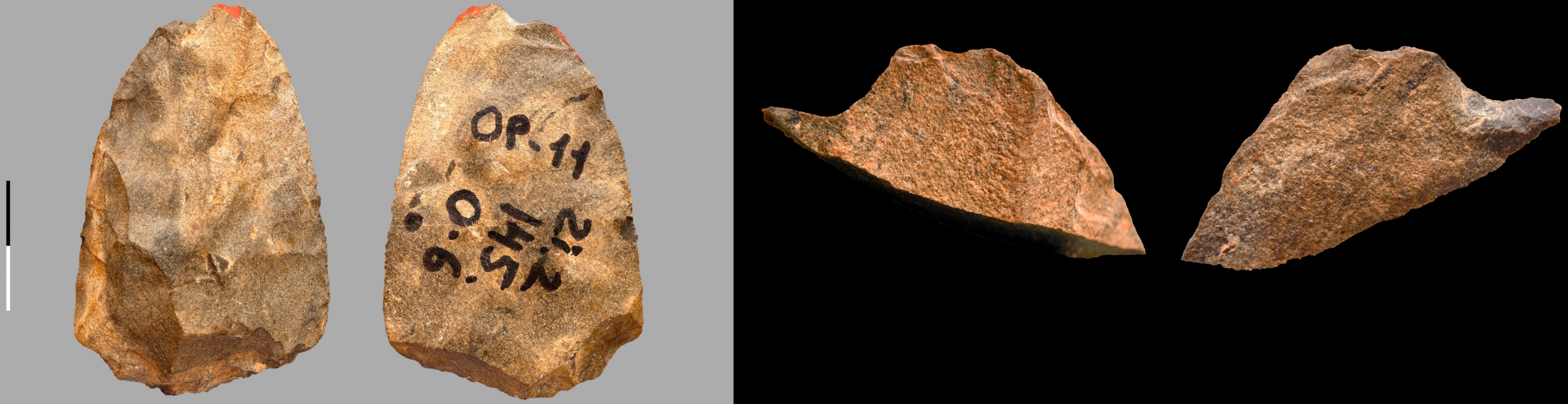


28 - OP 11 21. 2 06 KB 145

S3 Fig

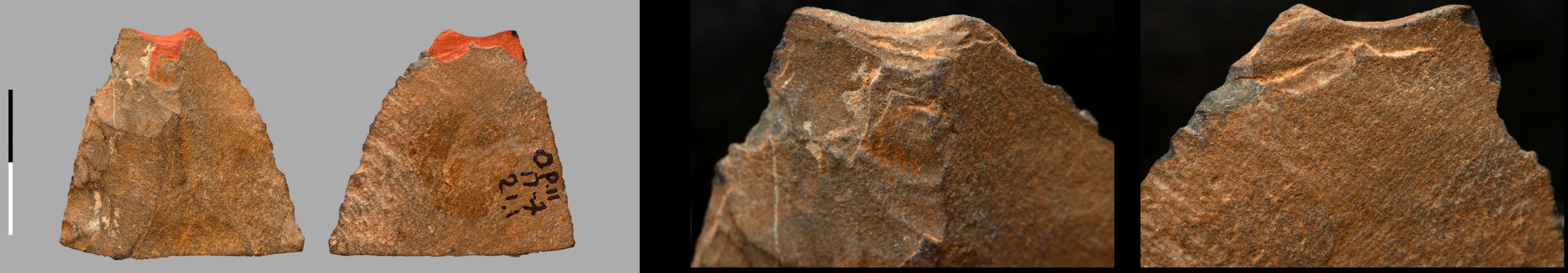


22 - OP 11 21.1

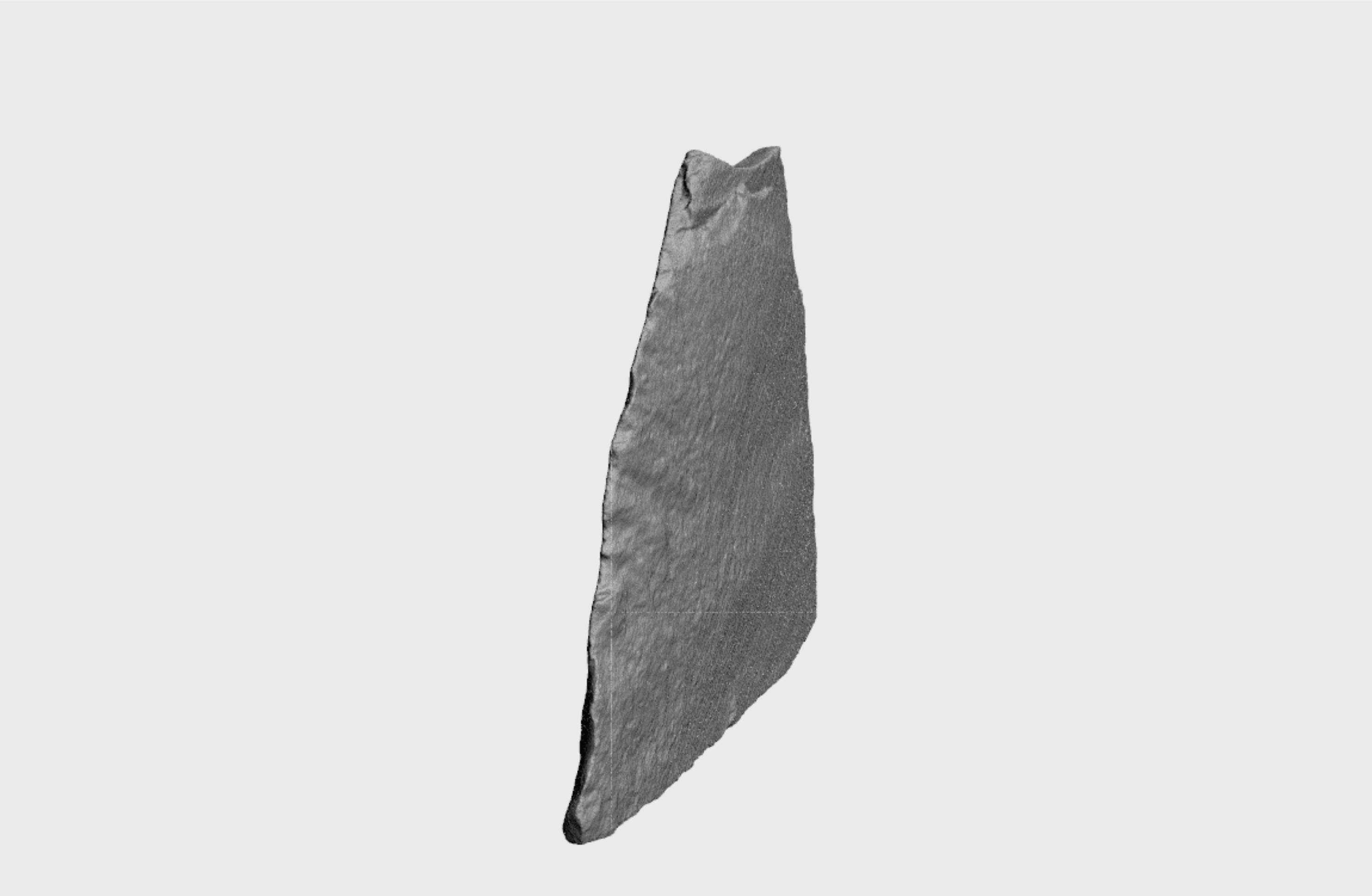

S3 File

Obi-Rakhmat: OP 11 21.1

S4 Fig

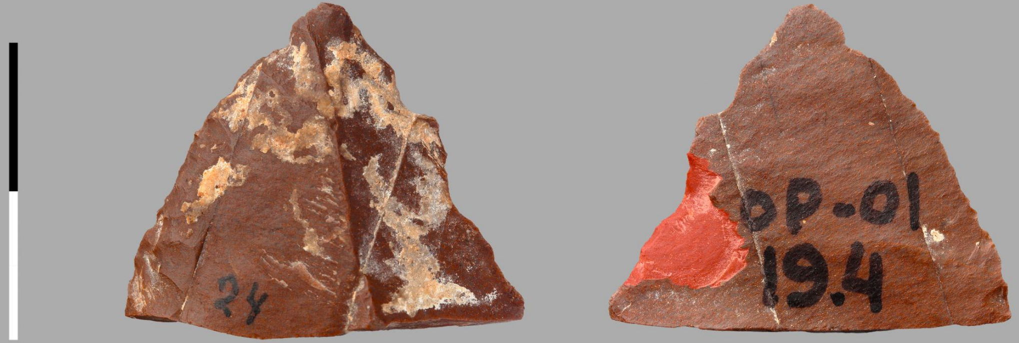


104 - OP 01 19 4

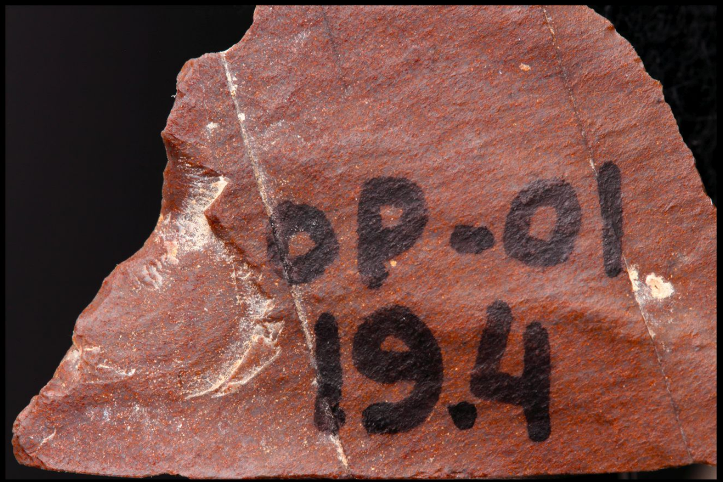

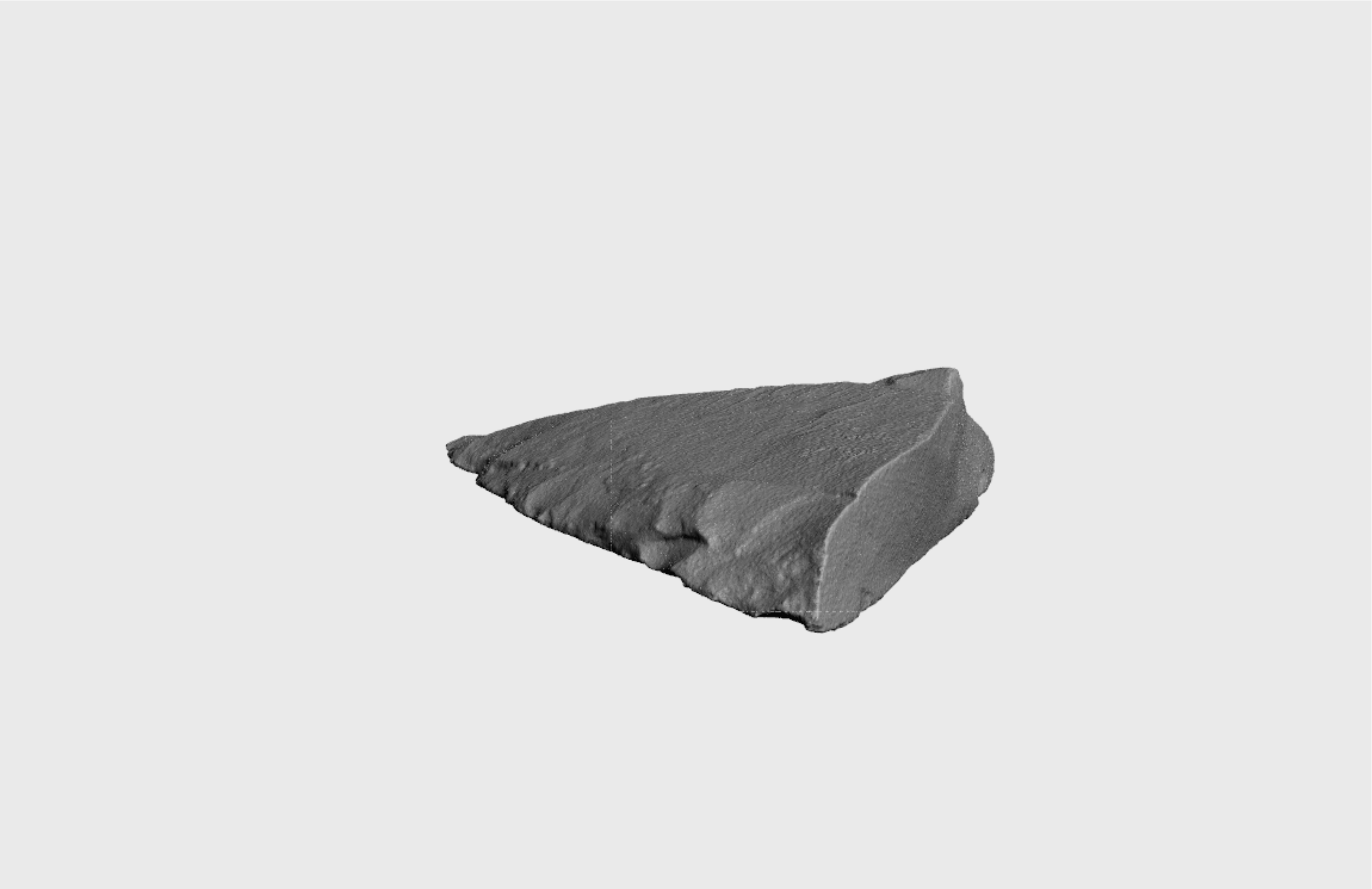

S4 File

Obi-Rakhmat: OP 01 19 4

S5 Fig

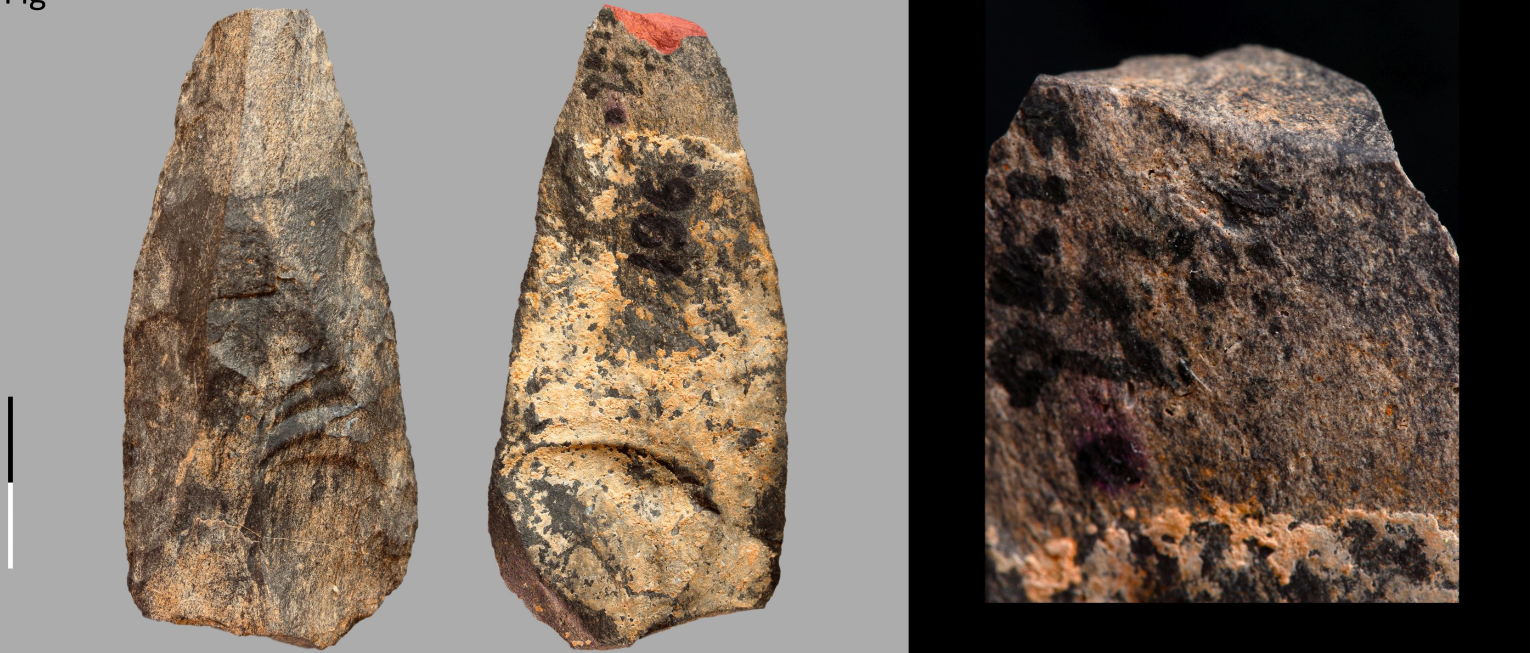

24 - ОР СЛ 21.1 196

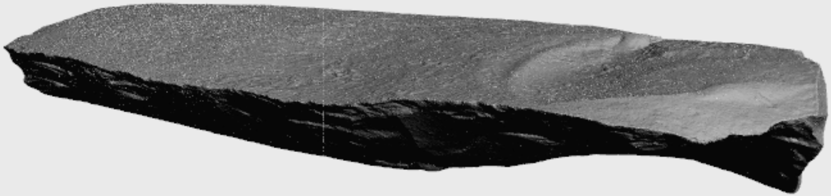

S5 File

Obi-Rakhmat: ОР СЛ 21.1 196

S6 Fig

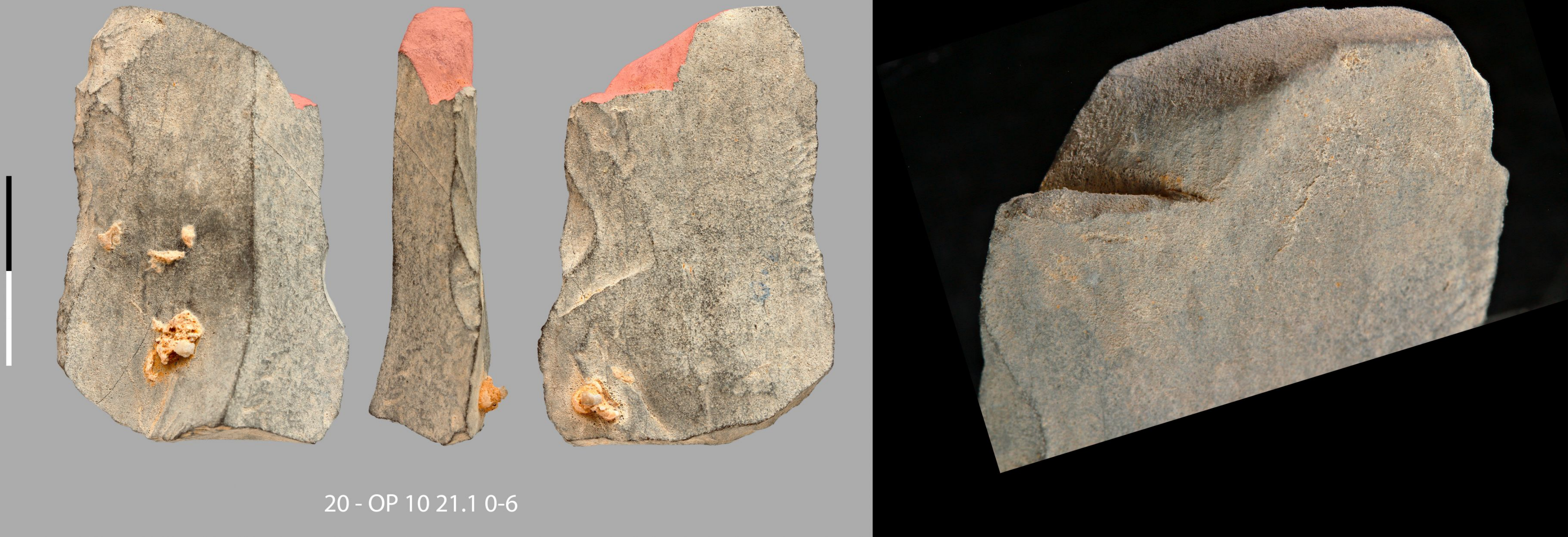

S7 Fig

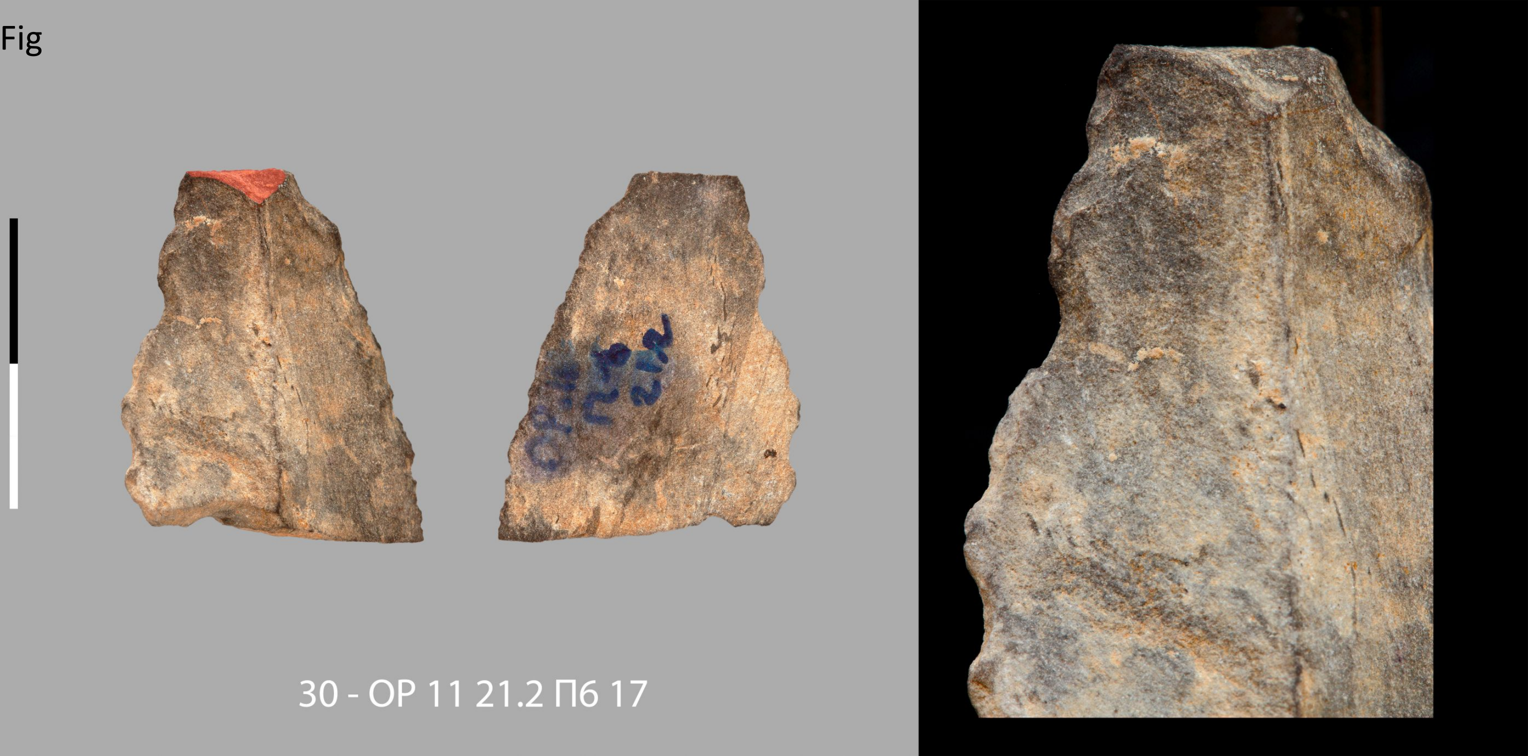

30 - OP 11 21.2 П6 17

S8 Fig

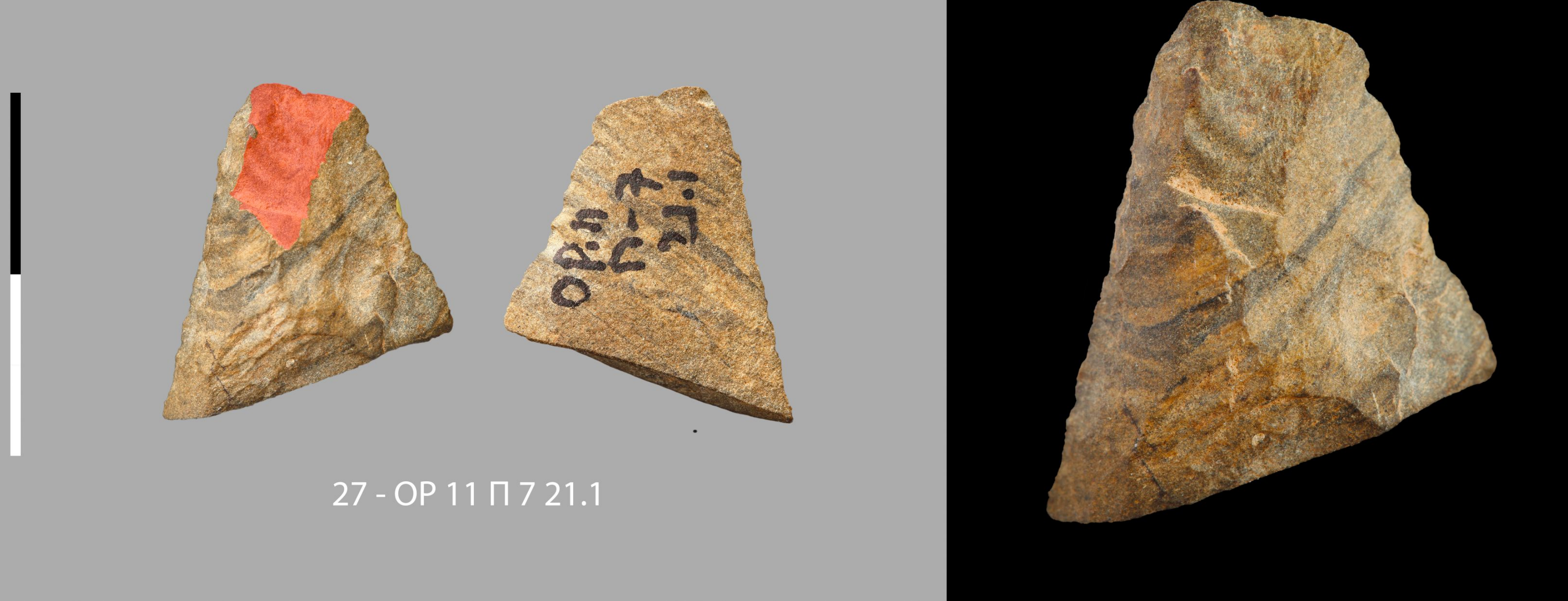

27 - OP 11 П 7 21.1

S9 Fig

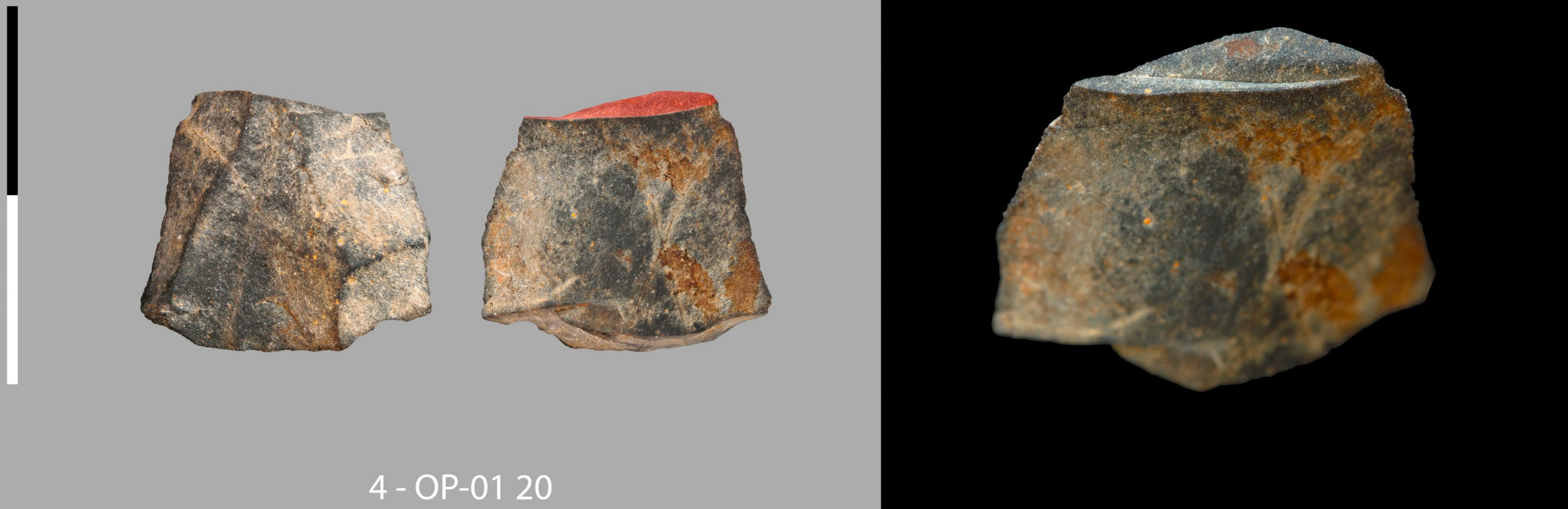

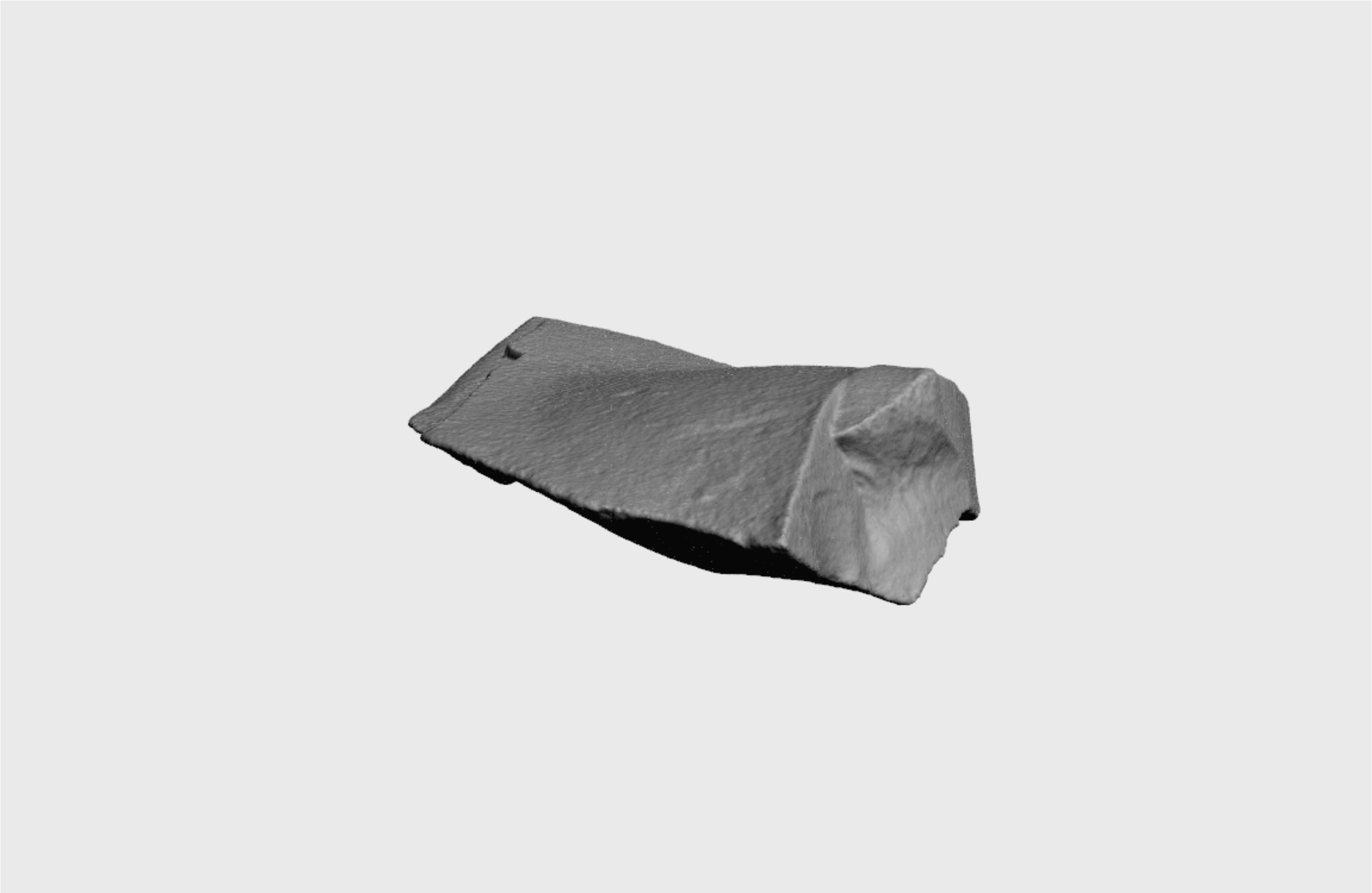

S9 File

Obi-Rakhmat: OP-01 20

S10 Fig

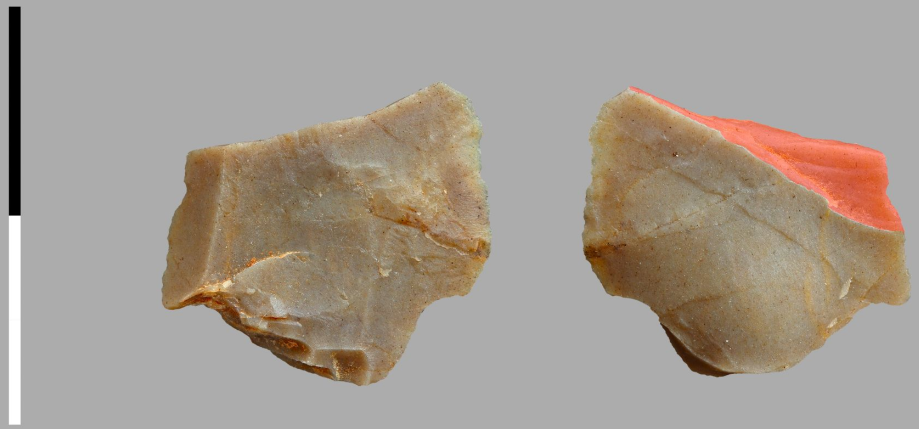

5 - OP 01-08 СЛ 21.1

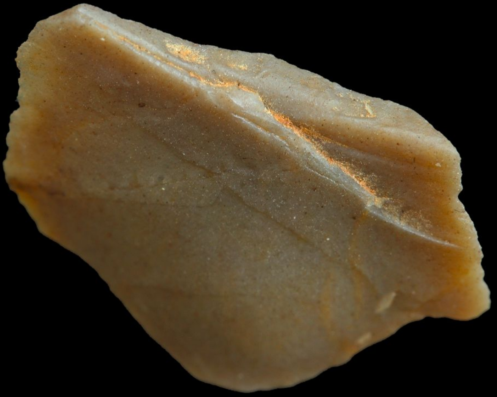

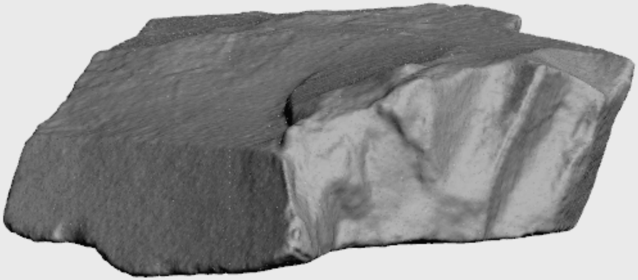

S10 File

Obi-Rakhmat: OP 01-08 СЛ 21.1

S11 Fig

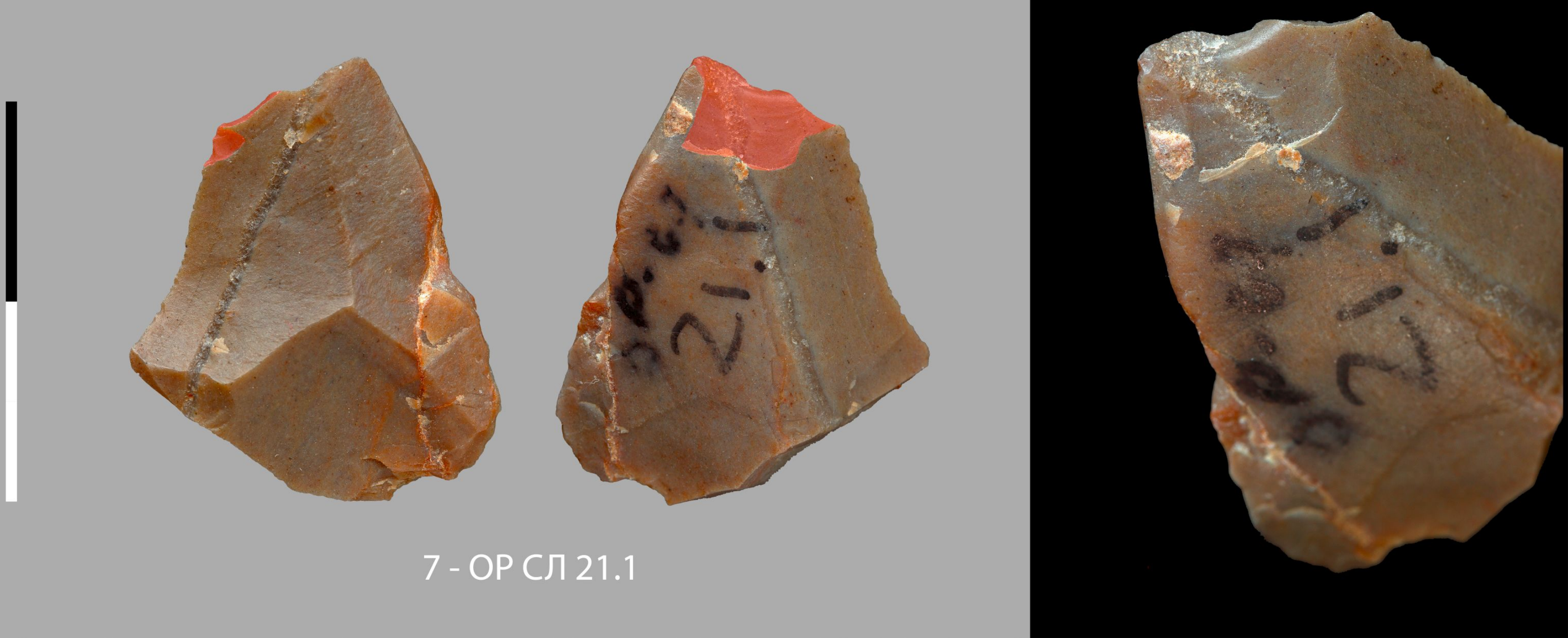

7 - ОР СЛ 21.1

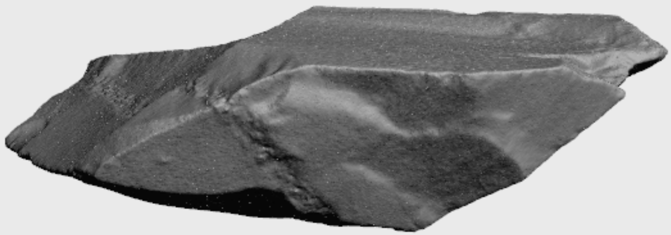

S11 File

Obi-Rakhmat: OP СЛ 21.1

S12 Fig

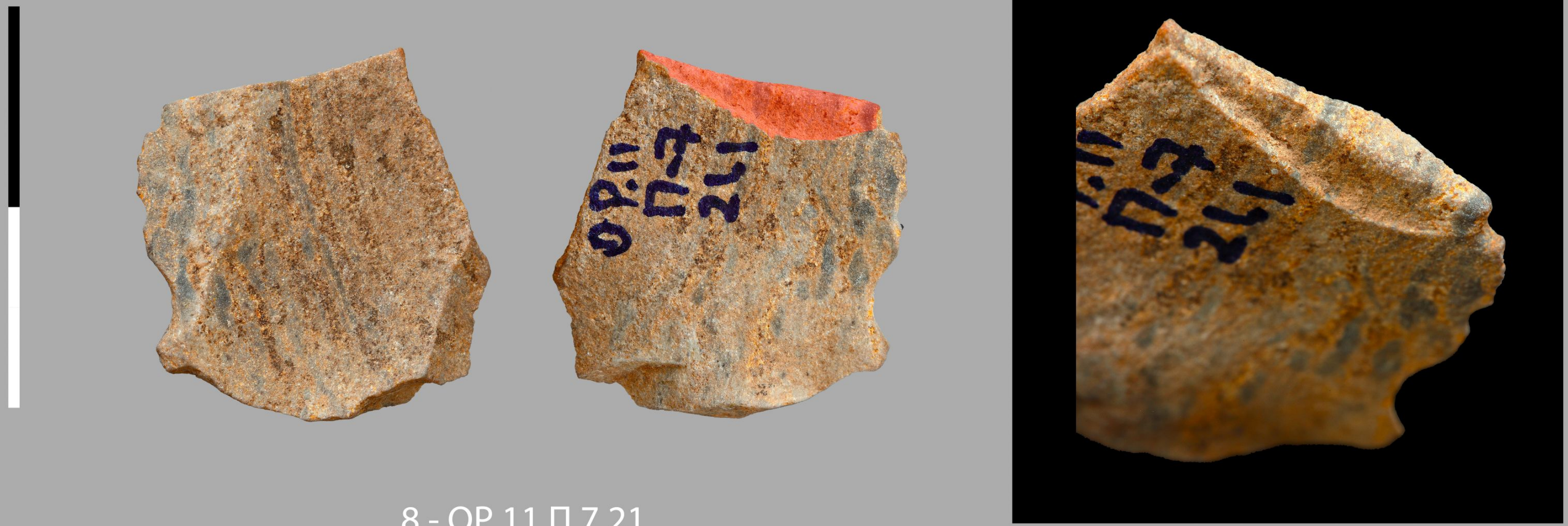


8 - OP 11 Π 7 21

S13 Fig.

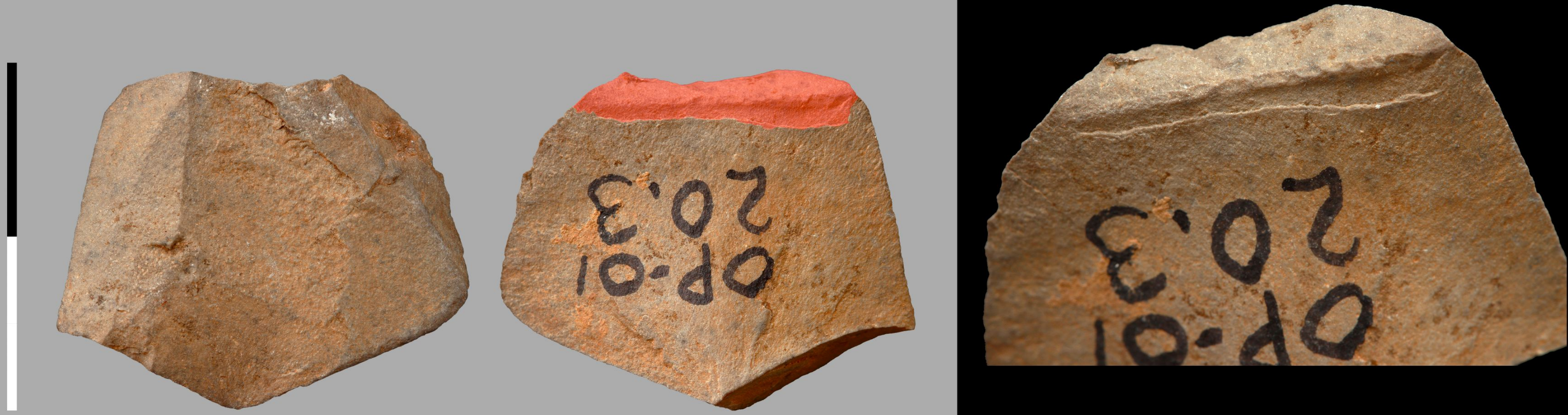

9 - OP 01 СЛ20.3

S14 Fig

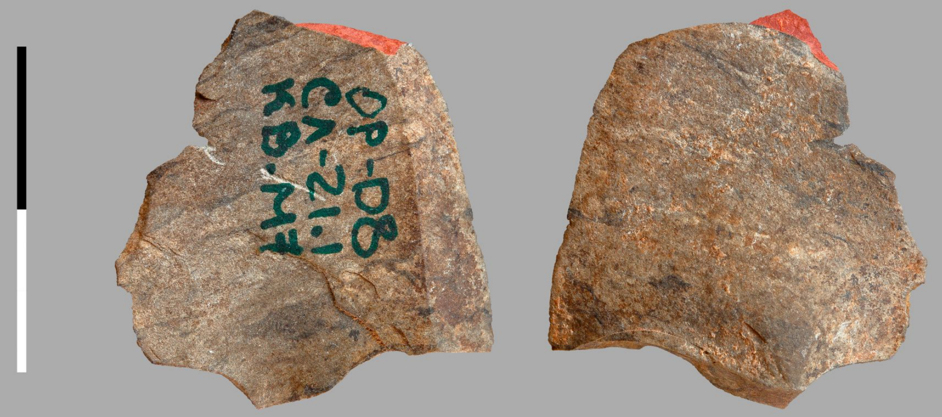

11 - OP 2001-2008 СЛ 21.1

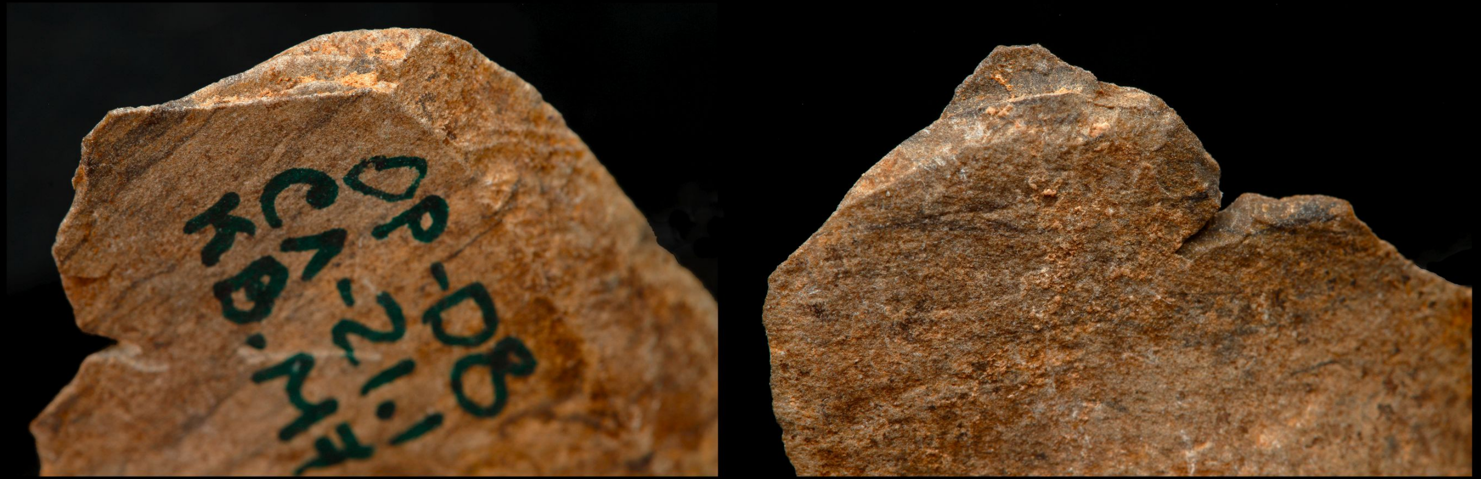

S15 Fig

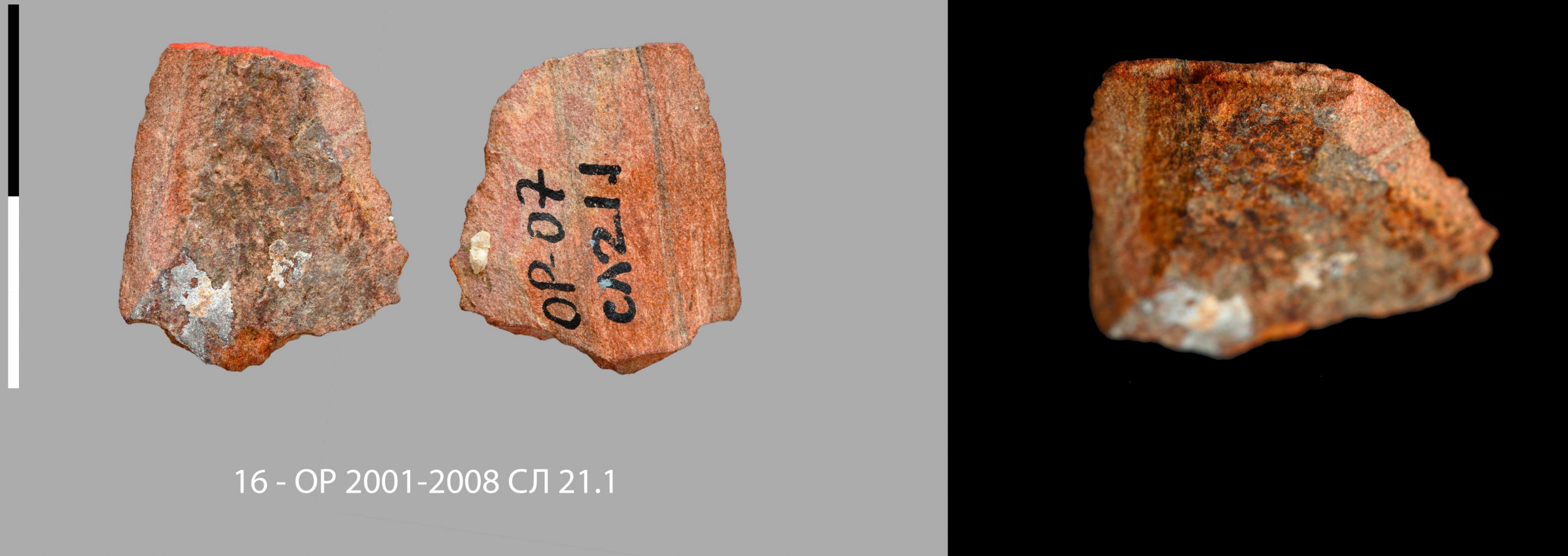


16 - OP 2001-2008 СЛ 21.1

S16 Fig

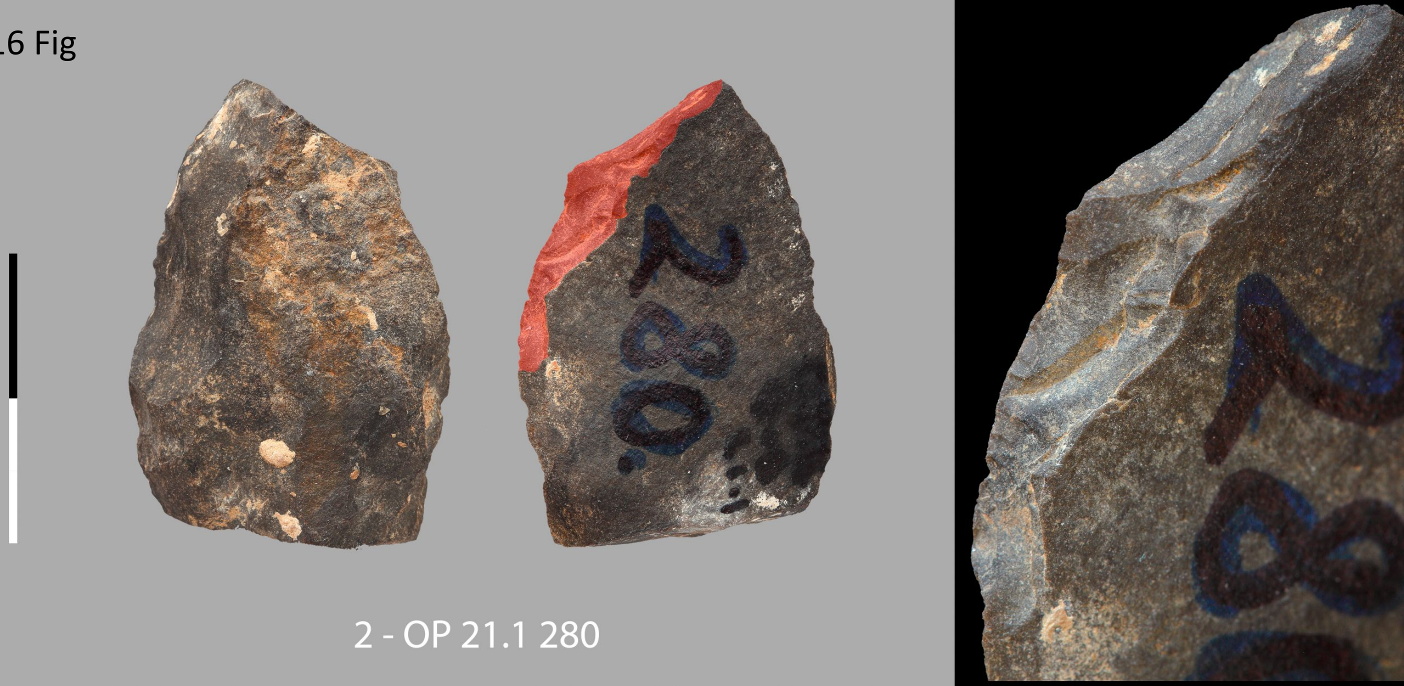

S17 Fig

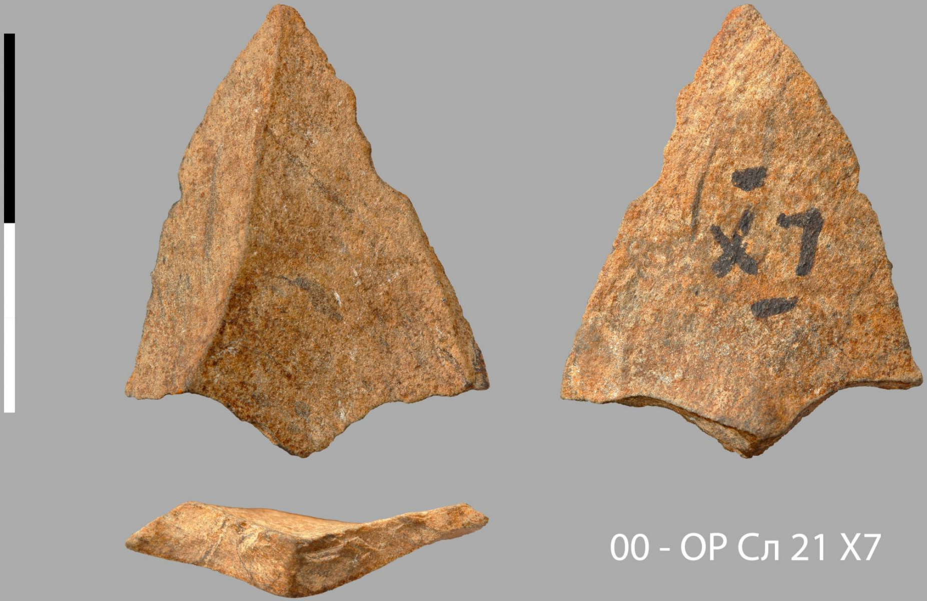

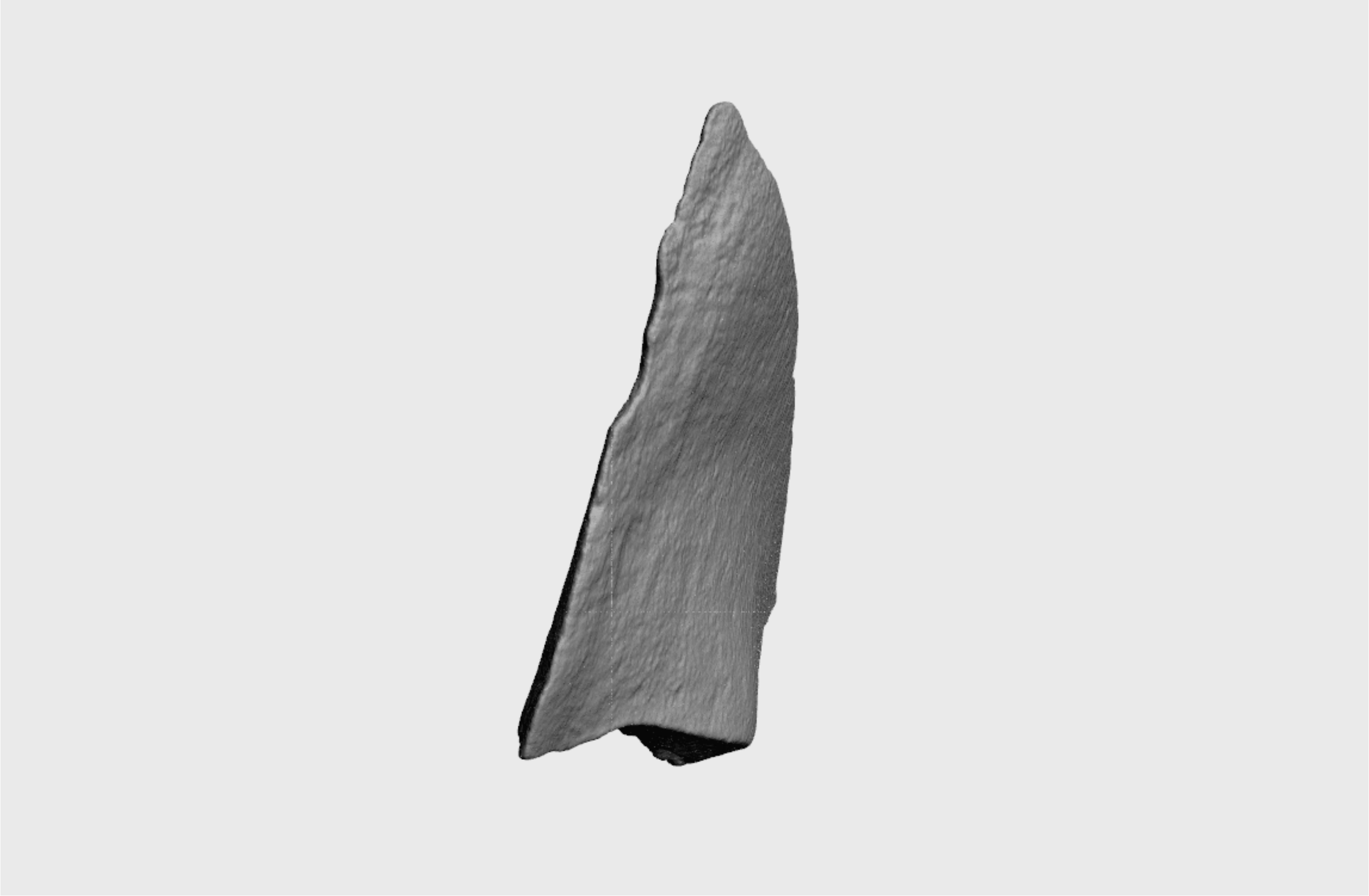

S17 File

Obi-Rakhmat: OP-10 21.1 H6 141

S18 Fig

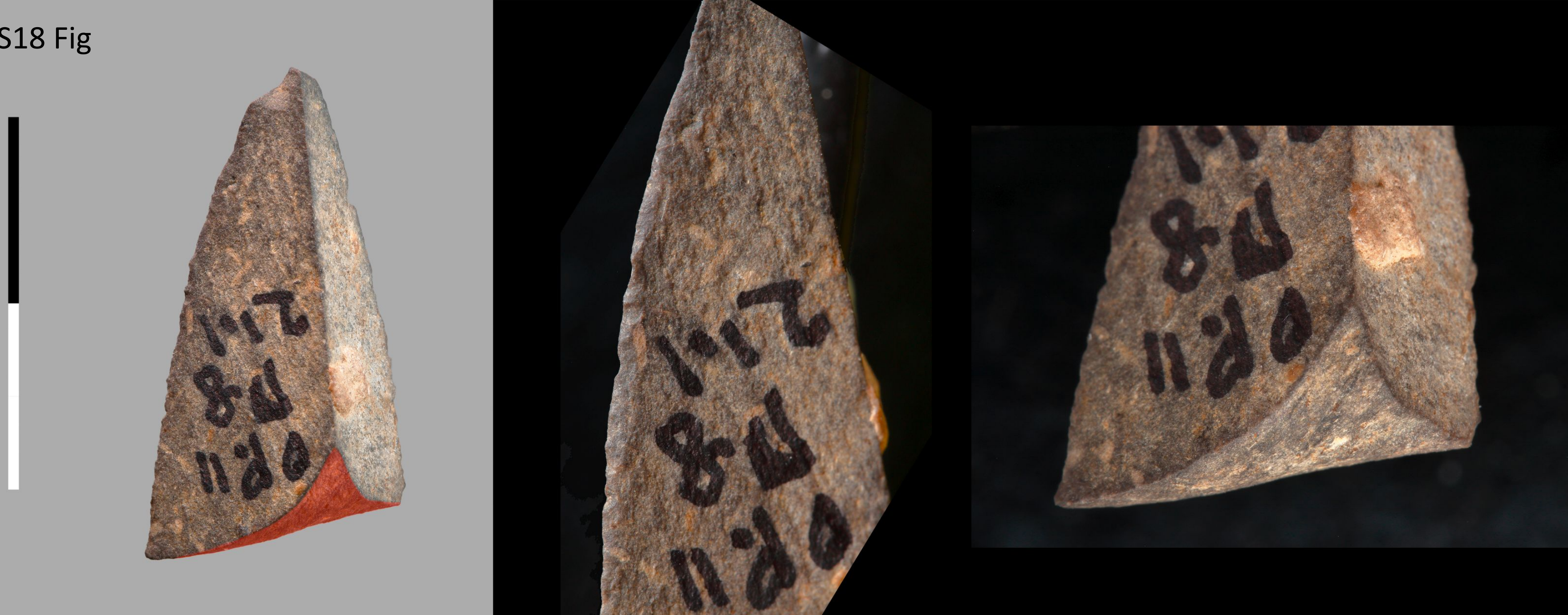

19 - ОР 11 СЛ 21.1 П8

S19 Fig

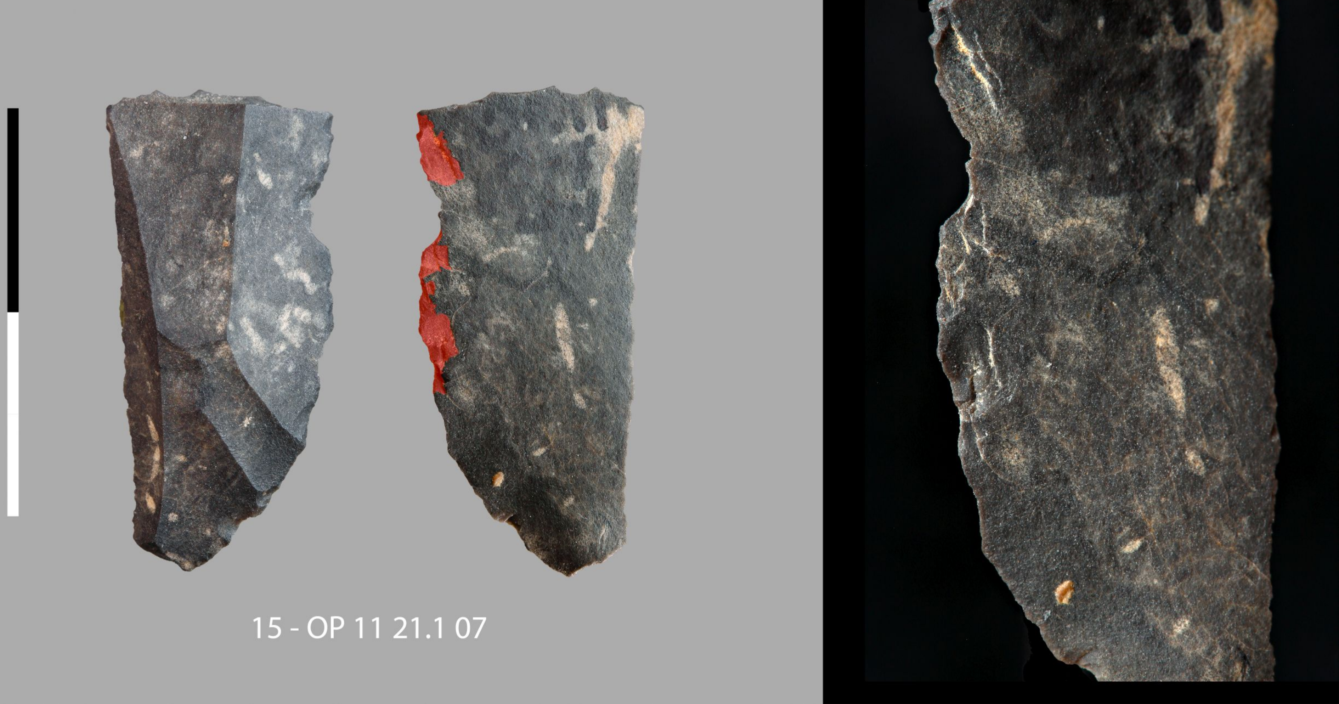

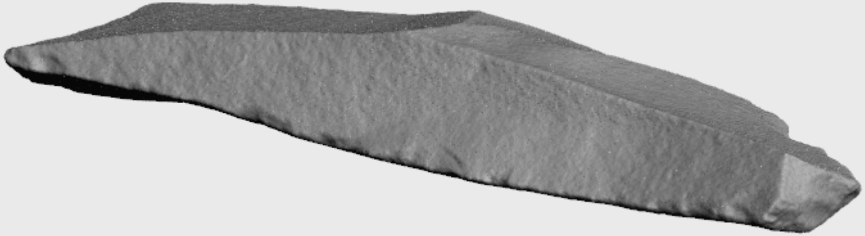

S19 File

Obi-Rakhmat: OP 11 21.1 07

S20 Fig

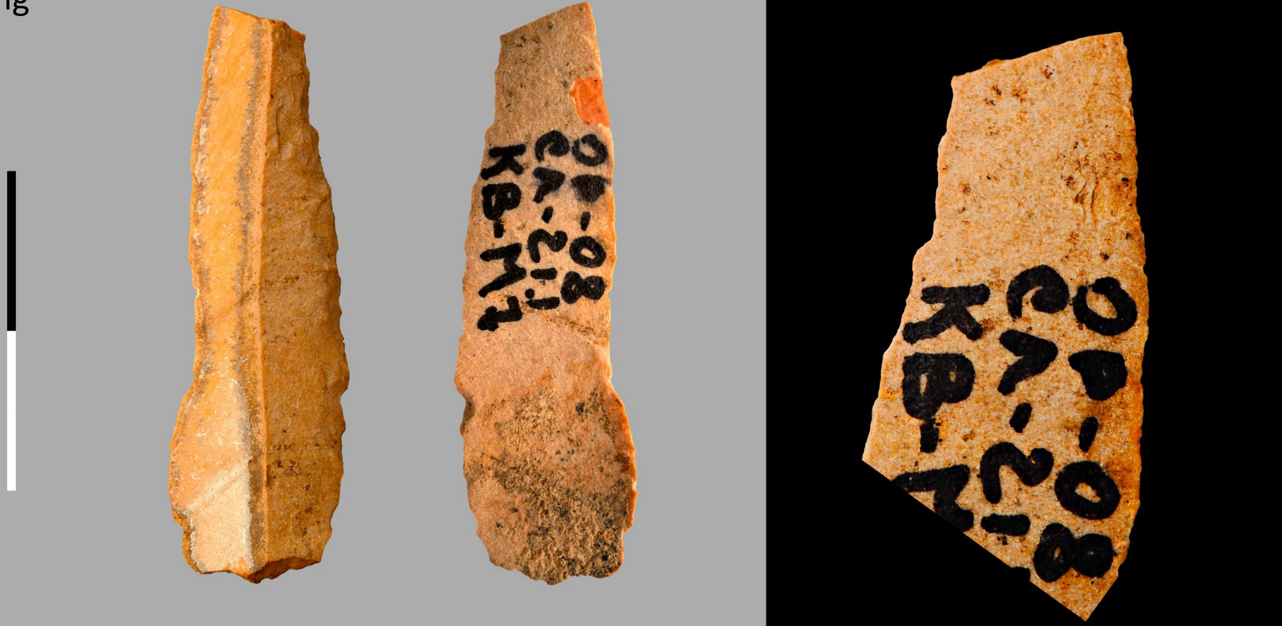

31 - ОП-08 Сл 21.1 КВ М7

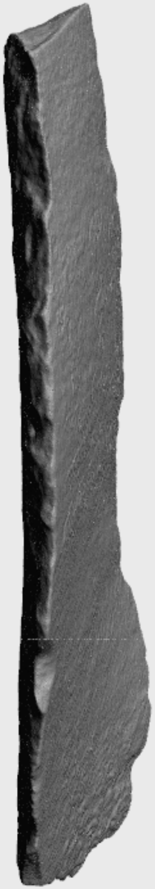

S20 File

Obi-Rakhmat: OP-08 Сл 21.1 КВ M7

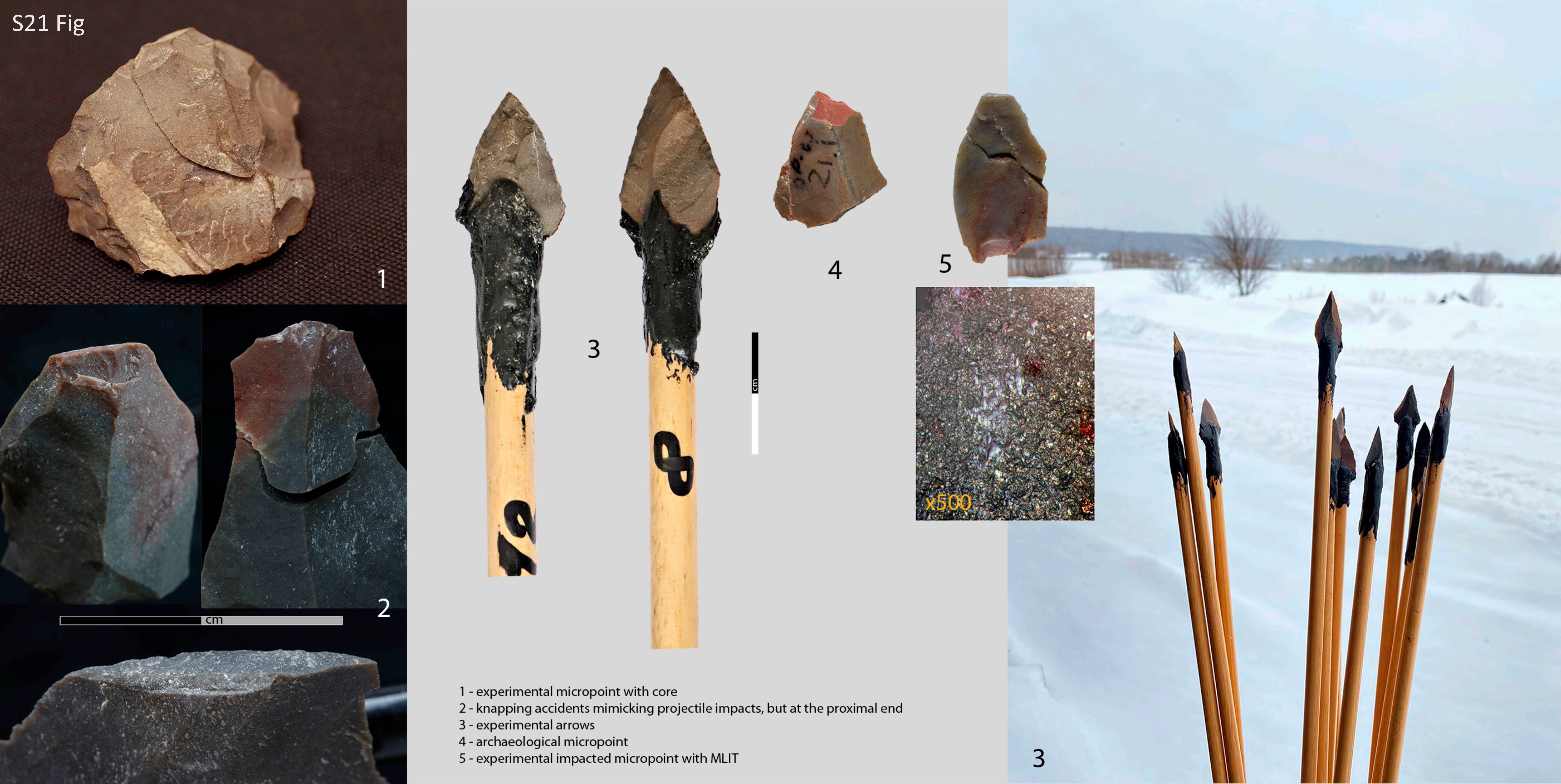


S21 Fig

1 - experimental micropoint with core
2 - knapping accidents mimicking projectile impacts, but at the proximal end
3 - experimental arrows
4 - archaeological micropoint
5 - experimental impacted micropoint with MLIT
